# Mitogen-activated protein kinase activity drives cell trajectories in colorectal cancer

**DOI:** 10.1101/2020.01.10.901579

**Authors:** Florian Uhlitz, Philip Bischoff, Stefan Peidli, Anja Sieber, Benedikt Obermayer, Eric Blanc, Alexandra Trinks, Mareen Lüthen, Yana Ruchiy, Thomas Sell, Soulafa Mamlouk, Roberto Arsie, Tzu-Ting Wei, Kathleen Klotz-Noack, Roland F Schwarz, Birgit Sawitzki, Carsten Kamphues, Dieter Beule, Markus Landthaler, Christine Sers, David Horst, Nils Blüthgen, Markus Morkel

**Author notes:** joint first authors (Florian Uhlitz, Philip Bischoff, Stefan Peidli).

## Abstract

In colorectal cancer, oncogenic mutations transform a hierarchically organized and homeostatic epithelium into invasive cancer tissue lacking visible organization. We sought to define colorectal cancer cell types and signals controlling their development. More than 30,000 epithelial single cell transcriptomes of tumors and matched non-cancerous tissues of twelve colorectal cancer patients were clustered into six patient-overarching groups defined by differential activities of oncogenic signaling pathways such as mitogen-activated protein kinase and oncogenic traits such as replication stress. RNA metabolic labeling and assessment of RNA velocity in patient-derived organoids revealed developmental trajectories of colorectal cancer cells organized along a mitogen-activated protein kinase activity gradient. This was in contrast to normal colon organoid cells developing along graded Wnt activity. Experimental targeting of EGFR-BRAF-MEK in cancer organoids affected signaling and gene expression contingent on predictive KRAS/BRAF mutations and induced cell plasticity overriding default developmental trajectories, providing a basis for non-genetic resistance to targeted therapies.

## Introduction

Healthy cells in the human body develop along trajectories controlled by intrinsic and extrinsic signals to ensure tissue homeostasis. Cancer cells cannot maintain homeostasis, as oncogenic mutations activate signaling pathways cell-intrinsically and render cancer cells unresponsive to paracrine signals (Hanahan & Weinberg, 2011). Colorectal cancer (CRC) commonly initiates via mutations activating Wnt/β-catenin signaling that maintains stem cells in the normal colon epithelium (Fearon, 2011). Subsequent mutations deregulate further signaling pathways such as RAS-RAF-MEK-ERK (also known as mitogen-activated protein kinase; MAPK signaling) providing pro-proliferative cues. Less frequently, CRC initiates via BRAF mutations, or from chronic inflammation increasing the mutation rate in the tissue (De Palma *et al*, 2019; Lasry *et al*, 2016). Genetic CRC drivers have direct and indirect effects on cancer cell development and the cellular composition of CRC and its microenvironment.

There is substantial evidence for the existence of tumor cell subpopulations and clonal architecture in CRC depending on genetics, microenvironmental cues, and space constraints (Van Der Heijden *et al*, 2019). Cancer stem cells are defined by their capacity for self-renewal and ability to initiate clonal outgrowth (Kreso & Dick, 2014). CRC cells with these characteristics have been distinguished by surface proteins like CD133, EPHB2 or LGR5 (Ricci-Vitiani *et al*, 2007; O’Brien *et al*, 2007; Merlos-Suárez *et al*, 2011). Likewise, lineage tracing in CRC cancer models has revealed preferential outgrowth of cancer cell subpopulations defined by expression of genes such as *LGR5* or *IL17RB* (Goto *et al*, 2019; Shimokawa *et al*, 2017) or by localization at the leading edge of the tumor (Lamprecht *et al*, 2017). Furthermore, it has been shown that CRC cells at the invasive front expressing genes such as the matrix metalloproteinase MMP7 contribute disproportionally to metastasis (Brabletz *et al*, 1999; Vermeulen *et al*, 2010). While these studies suggest that CRC cells are heterogeneous and hierarchically organized, other studies stress that developmental capacities of CRC cells are subject to a high degree of plasticity. In particular, oncogenic mutations and paracrine signals have been shown to trigger reversal of developmental trajectories so that differentiated cells regain stem cell characteristics (Schwitalla *et al*, 2013; Buczacki *et al*, 2013; Jadhav *et al*, 2017).

CRC is treated by chemotherapy and/or therapies targeting MAPK signaling, depending on predictive mutation status. Patients with metastatic CRC without RAS or RAF mutations profit from anti-EGFR antibody therapy (Van Cutsem *et al*, 2009; Karapetis *et al*, 2008), while patients with BRAF-mutant CRC now receive first-line therapy combinations of anti-EGFR antibodies and BRAF kinase inhibitors (Corcoran *et al*, 2018). Recent studies suggest roles for cell plasticity in therapy resistance. For instance, chemoresistance has been linked to subpopulations of CRC cells expressing the transcription factor ZEB2 (Francescangeli *et al*, 2020), and acquired anti-EGFR therapy resistance has been associated with stromal remodeling (Woolston *et al*, 2019).

Taken together, emerging evidence suggests that hierarchically organized tumor cell heterogeneity and cell plasticity play key roles in CRC progression and therapy response. However, developmental states of CRC cell are not well defined, and it is not known whether distinct differentiation states form preferential developmental trajectories. Here, we use single-cell RNA sequencing to identify patient-overarching CRC cell types defined by strengths of oncogenic signals and replicative responses. We use metabolic labelling of RNA in CRC organoids to delineate CRC development and show that CRC cell differentiation states, developmental trajectories and therapy-associated cell plasticity are informed by MAPK activity.

## Results

### CRC cells can assume distinct patient-overarching states

To capture the diversity of CRC cell states compared to the normal colon epithelium, we performed single-cell transcriptome analysis of twelve previously untreated CRC patients undergoing primary surgery (Fig. 1A). We utilized tissue samples that included the invasive tumor front and matched adjacent non-cancerous tissues (Supplementary Fig. 1). Tumors encompassed stages pTis (Tumor *in situ*) to pT4, with or without metastasis, and with various locations along the cephalocaudal axis of the colon (Supplementary table 1). Genetic analysis revealed mutational patterns characteristic for canonical CRC progression in most tumors; however, tumors from patients P007, P014, P020 and P026 contained the BRAF^V600E^ mutation often associated with the serrated progression pathway and tumor P008 was colitis-associated (Supplementary tables 1, 2). Eleven Patients were diagnosed with microsatellite-stable (MSS) CRC, while patient P026 was microsatellite-instable (MSI). We produced transcriptome libraries using a commercial droplet-based system and sequenced the libraries to obtain transcriptomes covering 500 to 5000 genes per cell. Transcriptomes were clustered and clusters were allocated to epithelial, immune or stromal subsets, using known marker genes (Smillie *et al*, 2019) (Fig. 1B, Supplementary Fig. 2, Supplementary table 3), and more than 30,000 epithelial cell transcriptomes were analyzed further.

**Figure 1:**
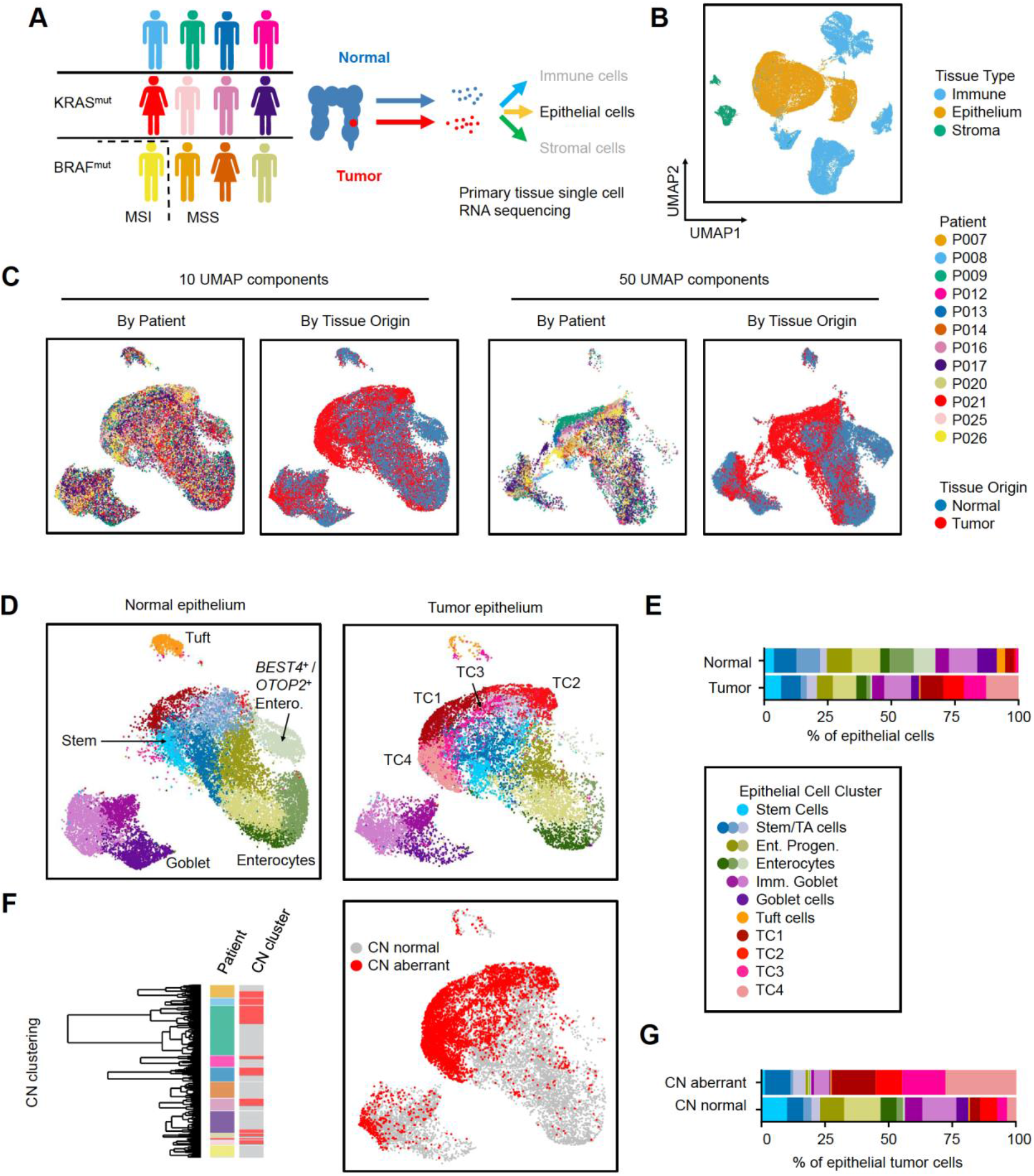
Generation of CRC single-cell RNA sequencing data and epithelial cell type census. **A** Workflow of the Clinical Single Cell Sequencing pipeline. In short, CRC and adjacent non-tumor tissue was sampled from 12 patients. Single cell RNA sequencing data was generated using the 10x Genomics platform, as outlined in Methods. For histology, see Supplementary Fig. 1. For patient characteristics, see Supplementary table 1, for mutational data, see Supplementary table 2. **B, C, D** UMAPs of single cell transcriptome data. **B** UMAPs of epithelial, immune, and stromal cell transcriptomes, color-coded by tissue origin as assessed by marker genes. For marker genes, see Supplementary table 3 **C** UMAPs of epithelial cell transcriptomes, color-coded by patient identity or tissue of origin, as indicated. D UMAPs of epithelial cells, separated by tissue of origin (normal vs. tumor). Clusters are color-coded by cell identity, as inferred from marker genes as outlined in main text. For epithelial cell cluster marker genes, see Supplementary table 4. **E** Relative fractions of epithelial cell types across all patients. For fractions per patient, see Supplementary Figure 3. **F** Identification of copy-number aberrant versus normal epithelial cells in tumor tissue. To the left: Cell cluster dendrogram, color-coded by patient and by copy-number-associated clusters (n=2 per patient). Copy-number normal cluster: grey; copy number-aberrant cluster: red. To the right: Distribution of SCN aberrant cells (red). For associated heatmap of gene expression and cell type data, see Supplementary Fig. 4. **G** Cell type assignment for SCN-normal versus -aberrant cells.

Epithelial cell transcriptomes of the different patients largely intermingled when visualized in a standard UMAP embedding employing ten principal components (McInnes *et al*, 2018), but partially separated when using a higher number of 50 components for UMAP embedding, particularly in areas enriched for tumor-derived transcriptomes (Fig. 1C). This suggests the existence of patient-specific gene expression patterns in cancer epithelium, but not in normal colon tissue. In summary, the UMAP embedding indicates that our single cell data are largely free from sample-specific bias, but instead reflect intrinsic differences between normal and tumor cell transcriptomes.

We used cell type-specific signatures and marker genes to annotate the epithelial cell clusters (Smillie *et al*, 2019; Fig. 1D, E, Figure EV1, Supplementary table 2). In the normal epithelium, we identified stem cells by markers such as *LGR5* and *OLFM4*. Neighboring clusters were annotated as enterocyte progenitors or mature enterocytes by expression of absorptive lineage markers such as *KRT20* and *FABP1* (Supplementary Fig. 3, Supplementary table 3). *BEST4*- and *OTOP2*-expressing enterocytes formed a discrete cluster (Parikh *et al*, 2019). Further separate epithelial clusters were identified as immature and mature secretory goblet cells expressing *MUC2* and *TFF3,* and as tuft cells expressing *TRPM5.* In tumor tissue, we additionally identified four tumor-specific clusters, termed TC1-TC4. These clusters were defined by high, but also unequal, levels of stem cell markers such as *OLFM4*, *CD44* and *EPHB2* and DNA repair genes such as *XRCC2*. *MMP7* was among the few genes expressed exclusively in the TC1-4 clusters, but not in the normal epithelium (Supplementary Fig. 3, Supplementary table 4). Clusters populated by differentiated absorptive and secretory cells were reduced in tumors, and profiles representing tuft cells and *BEST4*/*OTOP2*-positive enterocytes were vastly underrepresented.

MSS CRC is defined by somatic copy-number aberrations (SCNAs). Thus, we next distinguished cancer from non-cancer cells in the tumor tissues by inferring SCNAs from the single cell transcriptome data. We identified clusters of SCN-aberrant epithelial cells in ten out of the twelve tumors (Fig. 1F, Fig. EV2A). Exome sequencing of tumors P007, P008 and P009 validated SCNA calling from transcriptomes, showing that the procedure is robust for our single cell data (Fig. EV2B). P014 and P026 contained no cells with overt SCNAs. This was expected for tumor P026, which is MSI and thus defined by single nucleotide polymorphisms rather than SCNAs, but unexpected for P014, which was diagnosed as BRAF-mutant however MSS. In-depth analysis of patient-specific gene expression patterns (Supplementary Fig. 4A) and protein distributions (Supplementary Fig. 4B) revealed that patient-specific differences in cancer cell transcriptomes are at least partly driven by individual patterns of genomic gains and losses (Supplementary Fig. 4C-E, Supplementary table 5).

More than 86% of the TC1-4 cells were called SCN aberrant, along with substantial fractions of cells defined as stem cell/TA-like (36%) or immature goblet cells (22%, Fig. 1F, G). In contrast, a large majority (95%) of mature absorptive enterocytes and mature goblet cells derived from tumor samples were identified as copy number normal, and therefore likely stem from non-cancerous tissue at the tumor margins. In summary, our analyses define normal stem/TA-like cells, immature goblet-like cells, and TC1-4 cells as six main patient-overarching clusters of CRC cells.

### Epithelial tumor cell clusters differ by oncogenic traits and signals

We next defined characteristics of the six CRC cell clusters by assessing relative strengths of transcriptional footprints related to oncogenic signaling and cancer-associated functional traits (Fig. 2A, B). TC1 and TC4 were significantly enriched for the expression of direct ERK targets that are activated by MAPK (P=0.004 and P=0.006, FDR-corrected post-hoc Wilcox tests, respectively; Schubert *et al*, 2018), across all tumors and in particular in P007. We furthermore found a strong association of TC1 cells with the expression of hallmark signatures related to DNA repair and the G2/M replication checkpoint across all individual cancers (P=1.4E-10; Liberzon *et al*, 2015). This indicates that TC1 cluster cancer cells experience high levels of replication stress, a therapy-relevant trait of many cancers, including CRC. Indeed, TC1 cluster cells were exclusively assigned to the S or G2/M cell cycle phases by gene expression (Fig. 2C), in line with cells under replication stress, as also seen by *XRCC2* expression (Supplementary Fig. 3). The DNA damage-associated protein PARP stained many nuclei of the TC1-high CRC tissue P009 but not of TC1-low P008 (Fig. 2D). Cancer cells clustered in TC4 were characterized by expression of an intestinal YAP target signature (P=2.2E-8; Serra *et al*, 2019). YAP transcriptional activity is linked to regenerative responses and tumor progression (Zanconato *et al*, 2016). TC2 transcriptomes were significantly associated with high PI3K pathway activity (p=8.9E-10), related to control of metabolism and apoptosis. Wnt/β-catenin target gene activity was high across all TC clusters, but stem/TA-cell-like cancer cells showed stronger expression of a LGR5-ISC stem cell signature that is Wnt-driven (Merlos-Suárez *et al*, 2011; Muñoz *et al*, 2012), but this association was not significant across the patients. In summary, assessment of cell signaling signatures provides information on pathway activities of epithelial cancer cell clusters, and specific features of individual tumors. The analyses indicate that assignment of cancer cell transcriptomes to the TC1-4 clusters reflects, at least partially, differential states of oncogenic networks and oncogene-induced functional traits.

**Figure 2.**
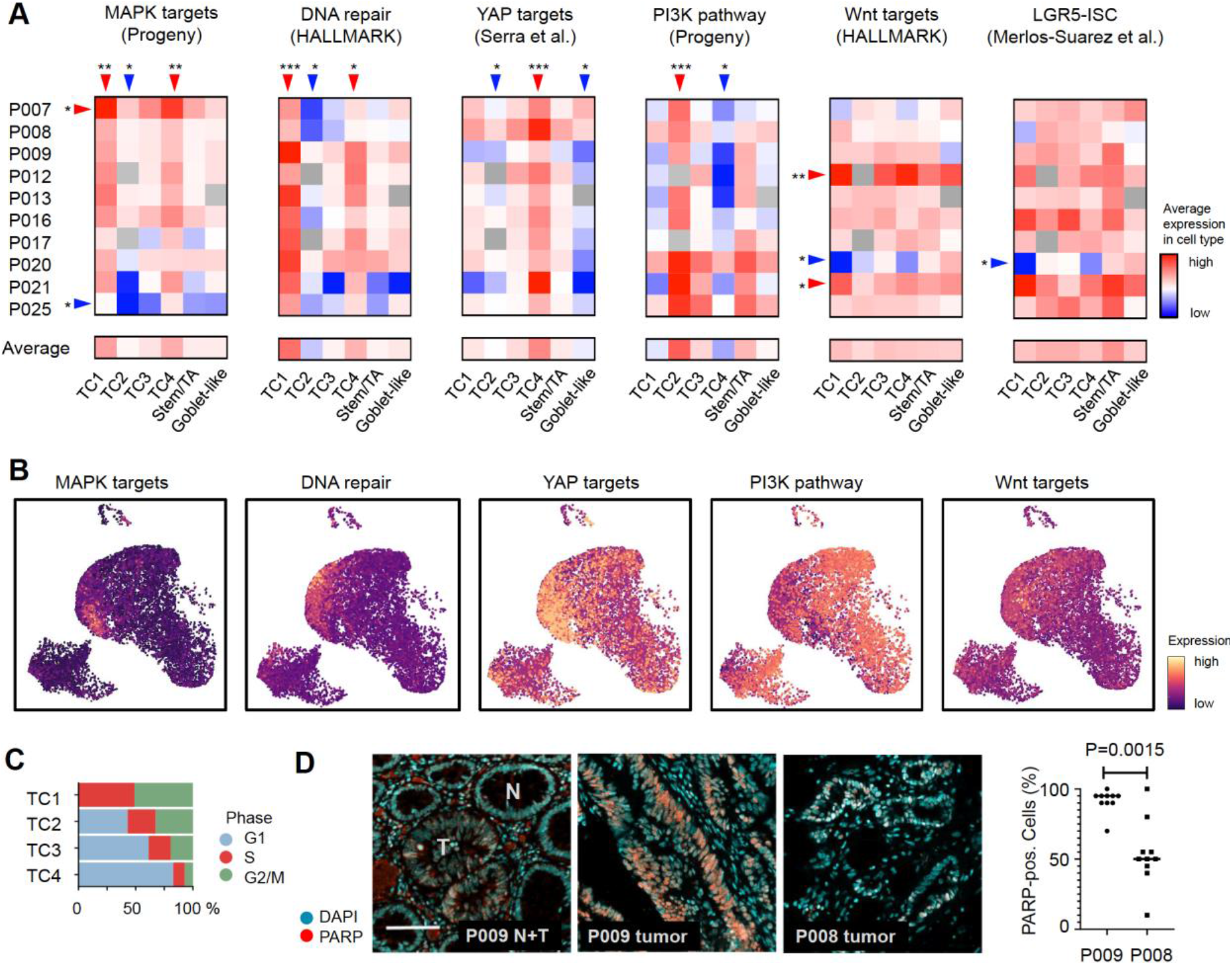
CRC cell clusters are distinguished by signaling pathway activities. **A** Transcriptional activity associated with key oncogenic traits and signals, by tumor-specific cell type and patient, as indicated. Red: High activity, blue: low activity. Significance was assessed by Kruskal Wallis Test (FDR corrected P<0.05), followed by a post-hoc analysis using a Wilcox-test of each group against all other groups, FDR corrected significance levels are shown (*=P<0.05; **=P<0.01; ***P<0.001). **B** Visualization of signatures corresponding to oncogenic traits and signals in the tumor cell transcriptome UMAP. **C** Cell cycle distribution of TC1-4 epithelial tumor cells, as inferred from single cell transcriptomes. **D** Immunofluorescence of DNA-damage-associated nuclear protein PARP. Images show adjacent normal and tumor crypts of tissue P009T, marked by N and T, respectively, and tumor tissue of patients P008 and P009, as indicated. Scale bar 100μm. Significance was assessed by an unpaired t-test, after blinded analysis of 10 random images per tumor.

### RNA metabolic labeling defines tumor cell hierarchies that correlate with MAPK activity

Immunofluorescent staining of primary CRC sections with antibodies directed against the stem cell marker OLFM4, the proliferation marker KI67 and the differentiation markers TFF3 and FABP1 revealed cell heterogeneity, but not how cancer cells in the tissue are related to each other (Fig. EV3). To establish whether CRC cells are hierarchically organized, we established organoid lines of two tumor samples, P009 and P013 (Fig. 3A). Organoids matched the cancer tissue on a mutational level (Supplementary Table 1). We cultured the cancer organoids, termed P009T and P013T, as well as normal colon organoids, termed NCO, in medium containing Wnt, R-Spondin, and EGF (WRE medium) or alternatively in medium lacking Wnt and R-Spondin (E medium). NCO organoids cultured in WRE medium showed graded expression of the intestinal LGR5-ISC stem cell signature, while expression of differentiation markers was graded in the opposite direction (Fig. 3B). LGR5-ISC signature activity was lost when NCO organoids were cultured in E medium. In P009T and P013T CRC organoids, LGR5-ISC signature activity was higher and independent of Wnt/R-Spondin, while expression of differentiation signature genes was much lower. Taken together, these expression patterns are in line with Wnt-dependent stem cell maintenance in normal tissue, and Wnt-independent stem cell maintenance and block of terminal differentiation in cancer tissue with APC mutations. The data however do not show whether graded developmental trajectories exist in CRC.

**Figure 3:**
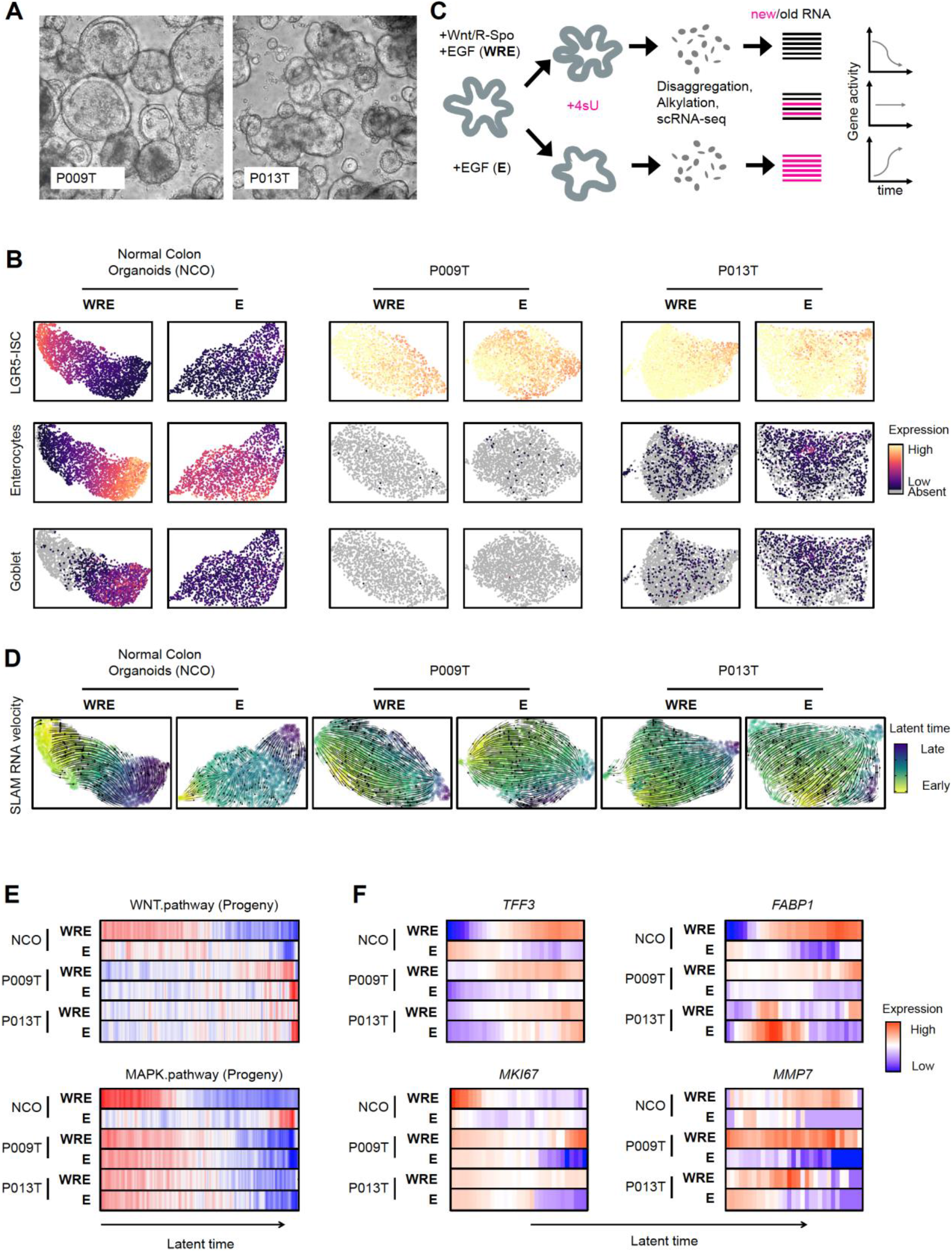
RNA metabolic labeling defines tumor cell trajectories in patient-derived organoids. **A** Phenotypes of patient-derived organoid lines P009T and P013T. **B** UMAPs of organoid single cell transcriptomes. Organoid lines and medium conditions as indicated. LGR5-ISC stem cell, enterocyte and Goblet cell signatures are visualized. **C** Schematic representation of SLAM-Seq workflow to infer RNA dynamics (“RNA velocity”). In short, organoids were passaged and assigned to different medium conditions. After three days, nascent RNA was metabolically labelled for 2 h using 4sU. Organoids were harvested, dissociated, and fixed. RNA in single cells was alkylated, and cells were subjected to single cell sequencing. Reads were assigned to nascent or old RNA status, depending on diagnostic T-C mutational status. **D** Developmental trajectories inferred from RNA metabolic labelling. Bold and thin arrows indicate high versus low directionality of RNA velocity. Colors below RNA velocity show latent time (yellow: early latent time; blue: late latent time). **E** Activities of Wnt/β-catenin and MAPK target genes in organoid single cell transcriptomes, ordered along latent time. **F** Activities of *TFF3*, *FABP1*, *MKI67* and *MMP7* in organoid single cell transcriptomes, ordered along latent time. Color code for panels D-F: Red: high activity; blue: low activity.

We, therefore, metabolically labelled RNAs of the organoids by 4-thio-uridine, before dissociation and single cell sequencing (scSLAM-Seq; Fig. 3C) (Herzog *et al*, 2017; Jürges *et al*, 2018). This allowed us to distinguish nascent labelled from older non-labelled mRNA, to order cells along inferred latent time based on dynamic RNA expression (Bergen *et al*, 2020), also known as RNA velocity (Supplementary Fig. 5 for quality controls). When cultured in WRE medium, developmental trajectories of normal NCO organoids initiated in areas of maximal LGR5-ISC signature scores and terminated in a region containing differentiated cells (Fig. 3D). When cultured without Wnt/R-Spondin in E medium, NCO normal colon organoids lost uniform direction of RNA velocity. In contrast, the P009T and P013T cancer organoids maintained strong transcriptional trajectories regardless of Wnt/R-Spondin in the medium.

We ordered organoid transcriptomes along latent time and assessed strengths of oncogenic signals (Fig. 3E). In line with the key role of Wnt in stem cell maintenance, normal colon organoids showed a gradient of Wnt/β-catenin target gene expression along latent time when cultured in WRE medium. In contrast, both P009T and P013T cancer organoids showed no graded Wnt/β-catenin-related expression, but a clear gradient of MAPK target gene activity along latent time in both medium conditions. *TFF3*, marking secretory differentiation, was graded in CRC organoids along latent time, but *FABP1*, marking absorptive differentiation in the normal colon, was not (Fig. 3F), in line with *TFF3* marking goblet-like CRC cells at the end of developmental trajectories. Proliferation marker *MKI67* was confined to the beginning of the latent time trajectory of normal organoids in WRE medium and showed extended gradients in CRC organoids. Both P009T and P013T organoids displayed Wnt-dependent loss of *MKI67* expression along latent time and also Wnt-dependent *MMP7* expression. In summary, our metabolic RNA labelling experiments indicate decreasing MAPK activity along CRC developmental trajectories and suggests a role for Wnt as a paracrine signal influencing gene expression, such as *MMP7*, in APC-deficient CRC cells.

### MAPK target gene expression defines CRC differentiation states

As MAPK-related gene expression was associated with developmental trajectories in CRC organoids, we analyzed whether the previously defined states of primary CRC cells are organized along a MAPK gradient in primary CRC. We assigned the SCN-aberrant primary cancer cell transcriptomes to 40 bins along a gradient of diminishing LGR5-ISC gene signature activity or along decreasing MAPK activity, and determined expression levels of the stem cell markers *LGR5* and *EPHB2* (Merlos-Suárez *et al*, 2011) (Fig. 4A). As expected, both *LGR5* and *EPHB2* were graded with LGR5-ISC signature strength; but expression of the MAPK target gene signature also ordered tumor cells in agreement with both CRC cell hierarchy markers and performed even better in sorting cells along a gradient of *EPHB2* expression. We conclude that expression patterns of known CRC cell hierarchy markers are in agreement with MAPK-driven development.

**Figure 4:**
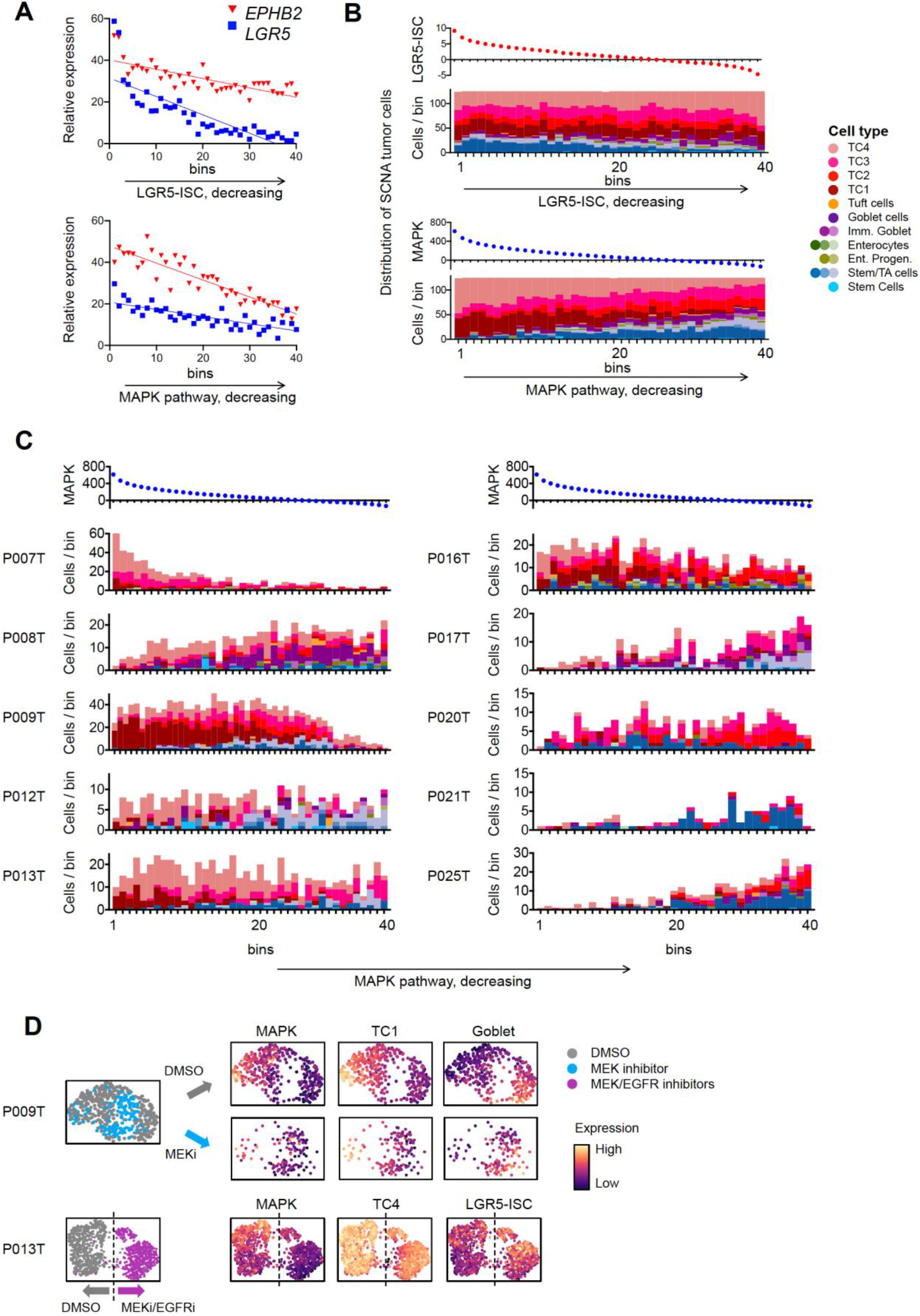
MAPK activity is linked to CRC cell differentiation states. **A** Gene expression of *LGR5* and *EPHB2*, along activity gradients of LGR5-ISC or MAPK target gene signatures. **B** Cell-type distribution of SCN-aberrant CRC cells along gradients of LGR5-ISC or MAPK transcriptional signatures, as in A. **C** Cell type distribution of SCN-aberrant CRC tumor cells along MAPK signature activity, as in B, per tumor. Cell fractions are color-coded by cell identity, as in Fig. 1C. **D** UMAP representations of single cell transcriptomes derived from P009T or P013T organoids, after MAPK blockade using MEK or combined MEK and EGFR inhibition. Color codes are treatment conditions or signature strengths, as indicated. Dashed line in P013T UMAP roughly separates control (DMSO) and MEK/EGFR inhibitor treated cells.

When ordering primary CRC transcriptomes along a gradient of LGR5-ISC activity, a higher proportion of stem/TA-like tumor cells aggregated at the high end of the gradient, whereas tumor cells assigned as immature goblet cell-like accumulated in the lower end, and TC1-4 cells displayed a broad distribution (Fig. 4B). In contrast, ordering of CRC cells by MAPK signature activity significantly enriched TC1 and TC4 cells at the start of the gradient (P=8.6E-23 and P=7.6E-20, respectively, adjusted Pearson’s chi-squared p-value), whereas stem/TA- and immature goblet cell-like cells concentrated at the lower end (P=1.3E-19 and P=1.1E-8, respectively). These aggregate differences were also reflected in cell state distributions of individual patients (Fig. 4C): P007 showed the highest MAPK activity, and the highest proportion of TC4 cells which clustered disproportionally in the MAPK-high bins. TC1 and TC4 cells had also the highest average MAPK activities in most other CRCs, including P008, P012, P013, and P016. In contrast, stem-/TA-like tumor cells and immature goblet-like CRC cells displayed relatively low expression of MAPK targets in all tumors, particularly in P009, P012, P013 P017, P021, P025 and P008, P017, respectively.

To experimentally determine whether the CRC cell states are functionally linked to MAPK activity, we blocked MAPK signaling in CRC organoids by the MEK1/2 inhibitor Selumetinib (AZD6244) or by Selumetinib in combination with the EGFR inhibitor Sapatinib (AZD8931) and analyzed singe cell gene expression after 48h (Fig. 4D). We found that cells showing a high TC1 signature were diminished after MEK inhibition in P009T organoids, while the fraction of cells with a high goblet-like signature was increased. Likewise, P013T organoid cells showed lower expression of the TC4 signature genes after MAPK blockade, and higher expression of the stem cell-related LGR5-ISC signature. These results suggest that CRC cell states are MAPK-driven.

### Targeted therapy alters cell signaling networks and differentiation signatures in CRC organoids

MAPK is a key target of CRC therapy, as blockade of EGFR is the first-line therapy for patients with metastasized MAPK-wildtype CRC, and combined blockade of EGFR and BRAF is first-line therapy for patients with advanced CRC containing oncogenic BRAF mutations. We therefore asked whether targeted therapy is associated with changes in CRC cell states and trajectories. In addition to the RAS/RAF-wildtype organoid lines P009T and P013T, we employed OT227 and OT302 organoids carrying KRAS^G13D^ and KRAS^G12V^ mutations, respectively, and the BRAF^V600E^-mutant lines B2040 and C2019. As the clinically relevant blockade of EGFR using the antibody Cetuximab was not effective *in vitro* as also observed by others (Schütte *et al*, 2017), we treated the organoids with the EGFR inhibitor Sapatinib, the BRAF inhibitor LGX818 (Encorafenib), the MEK inhibitor Selumetinib and combinations for 48h, before subjecting single cell suspensions to CyTOF and scSLAM-seq (Fig. 5A, Supplementary Fig. 6 for summaries of CyTOF data) to measure relative activities of signal transducers and related transcriptional signatures, respectively (Fig. 5B).

**Figure 5:**
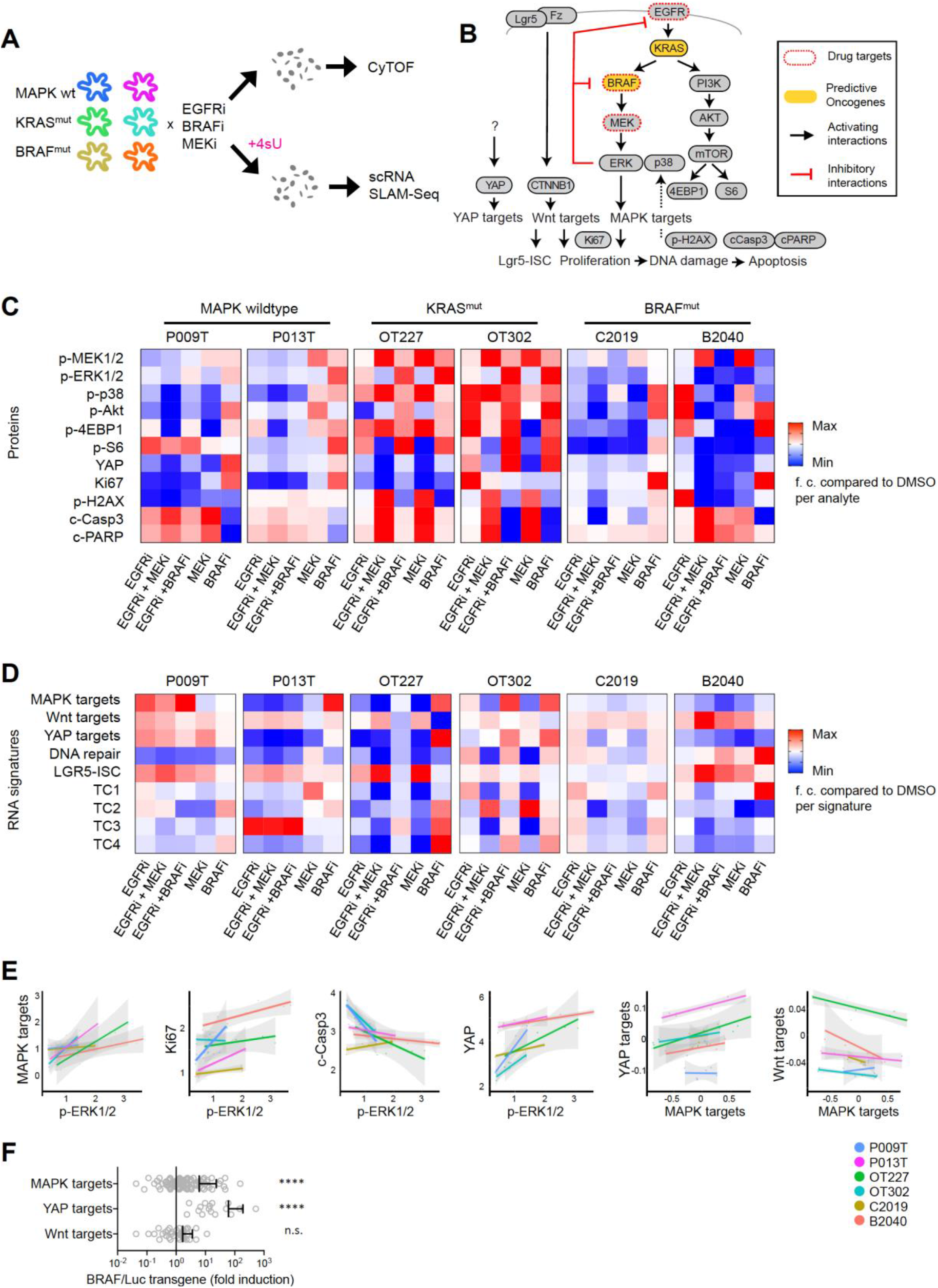
Anti-MAPK therapy affects signaling networks and transcriptomes contingent on predictive mutations in organoids. **A** Workflow of the experiment. In short, organoids were treated for 48h with inhibitors, before disaggregation into single cells for CyTOF and scRNA-seq analysis. For scRNA-seq, organoids were labelled for 2h with 4sU. **B** Schematic representation of signaling network and transcriptional signatures associated with phenotypes or signaling pathways, as indicated. **C** Heatmap of key CyTOF data. Average activities of selected analytes are given as log fold-change after normalization to DMSO control condition. Range of color scale was adjusted for each analyte. For relative changes between all analytes, see Supplementary Fig. 10). **D** Heatmap of key scRNA seq data. Average activities of selected gene signatures are given as log fold-change after normalization to DMSO control condition. Range of color scale was adjusted for each signature. **E** Correlation between and within the CyTOF and scRNAseq data sets. For each line, average protein analyte (CyTOF) or transcriptional signature (scRNA seq) values were plotted for all six experimental conditions. Graphs give trend line and confidence intervals. **F** Upregulation of MAPK and YAP target genes in mouse intestinal organoids after induction of oncogenic BRAF. Wnt target genes is not significantly affected. For experimental details, see Riemer *et al.* (2015).

The effects of inhibitor treatments on the signaling network were variable between lines, but we also observed communalities between the lines with shared RAS/RAF mutational status (Fig. 5C). In RAS/RAF-wildtype lines, treatment with EGFR inhibitor reduced both MEK and ERK phosphorylation. MEK inhibition decreased ERK phosphorylation, but increased MEK phosphorylation via negative feedback suppression (Fritsche-Guenther *et al*, 2011). BRAF inhibitors increased levels of MEK and ERK phosphorylation, suggesting paradoxical activation of RAF (Hatzivassiliou *et al*, 2010). In contrast, the KRAS-mutant lines OT227 and OT302 were largely unresponsive to EGFR inhibition, while MEK inhibition caused strong upregulation of MEK phosphorylation, and BRAF blockade strongly upregulated both MEK and ERK phosphorylation. A similar response to MEK inhibition was found in the BRAF-mutant lines C0219 and B2040, however in these lines BRAF inhibition alone or in combination with EGFR inhibition resulted in substantial loss of ERK phosphorylation. Across all lines, we observed a positive correlation between phosphorylation of ERK and the Ki67 (P<0.005, using linear models with line-specific offsets) and a negative correlation between p-ERK and cleaved Caspase3 (P<0.005) and cleaved PARP levels, in line with roles of MAPK in activation of proliferation and inhibition of apoptosis.

Furthermore, we found high correlation between ERK phosphorylation and expression of MAPK target genes (P<0.0001, using linear models with line-specific offsets; Fig. 5D, E), between p-ERK and YAP protein levels (P<0.0001), and between MAPK target expression and YAP target expression (P<0.0001). In contrast, MAPK target activity was negatively correlated with Wnt target activity (and P<0.05, respectively; Fig. 5E). A direct interaction between MAPK and YAP signaling is supported by induction of transgenic BRAF^V600E^ in a mouse intestinal organoid model (Riemer *et al*, 2015), which resulted not only in activation of the MAPK target gene signature, but also in an even stronger activation of YAP target genes (Fig. 5F). Our findings thus indicate that the MAPK, YAP and Wnt signaling pathways form an interconnected network controlling transcription, and ultimately proliferation and cell fate in CRC. Accordingly, we found that experimental inhibition of MAPK also modulated prevalence of cell-type-related signatures (Fig. 5D): in P013T, OT227 and B2040 organoid lines, effective repression of ERK phosphorylation was correlated with activation of the LGR5-ISC stem cell-related signature, while gene expression related to the MAPK-high cell states TC1 and TC4 was downregulated under these conditions in all lines.

### Targeted therapy alters developmental trajectories of CRC organoid cells

For a detailed analysis of CRC cell development under experimental MAPK therapy, we assessed transcriptome data on single cell level. Anti-MAPK therapy had broad consequences on gene expression, as transcriptomes derived from conditions of effective ERK suppression often inhabited different spaces in a UMAP representation (Fig. 6A), as quantified by hierarchical clustering of conditions based on a similarity matrix of shared neighboring cells (Fig. 6B). We found that inhibition of EGFR, alone or in combination with BRAF or MEK inhibition, was effective in the induction of transcriptome changes in MAPK-wildtype organoids. Likewise, a combination of BRAF and EGFR inhibitors deregulated transcriptomes effectively in the BRAF-mutant organoid B2040, while both inhibitors were not as effective on their own. BRAF-mutant C2019 organoids were largely unresponsive to anti-MAPK treatment. As also observed for ERK phosphorylation (Fig. 5C, above), these effects agree with clinical reality, where positive outcomes for EGFR and combinatorial BRAF/EGFR inhibition are variable but limited to patients with RAS/RAF wildtype and BRAF-mutant CRC, respectively.

**Figure 6:**
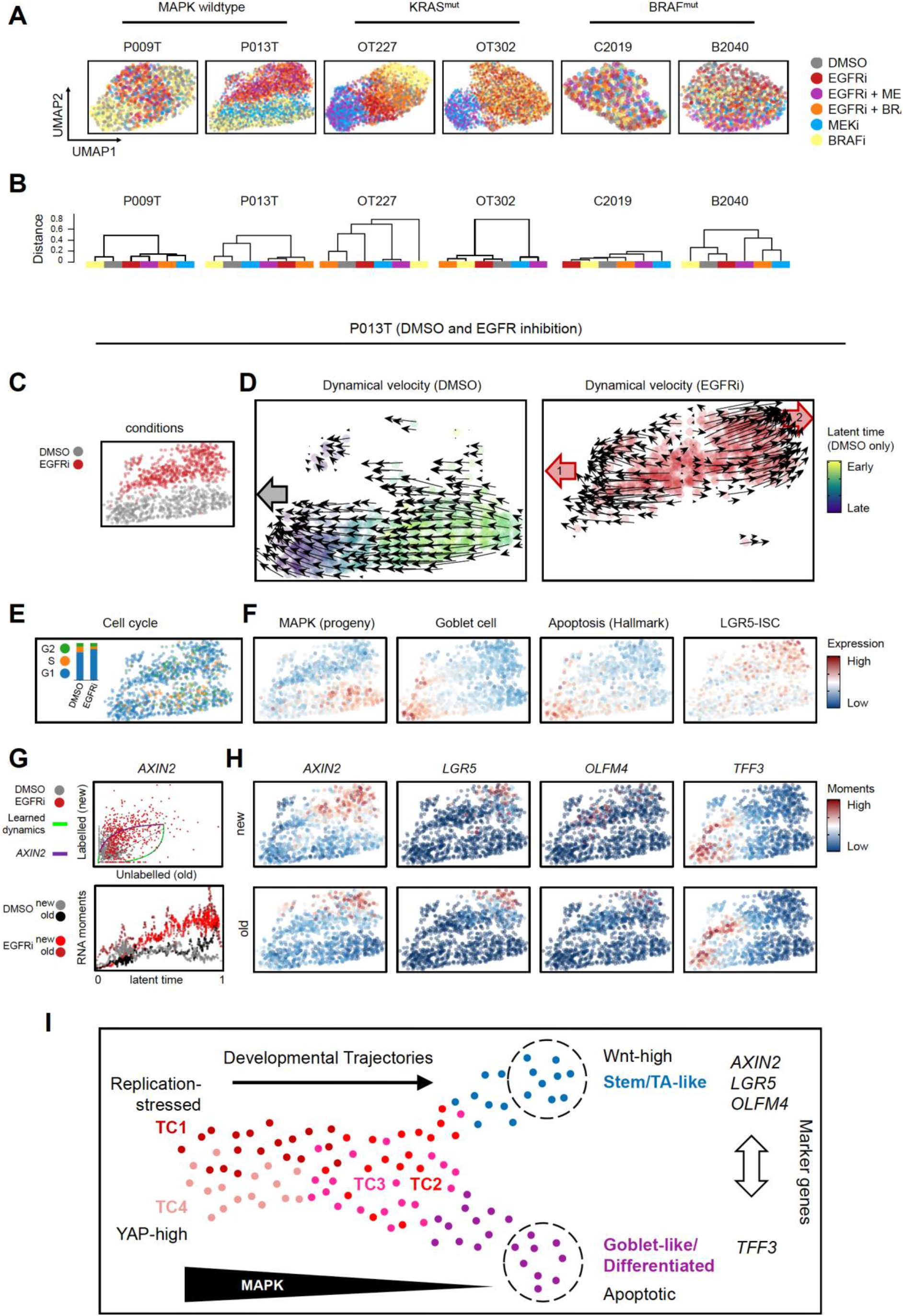
Anti-MAPK therapy re-routes developmental trajectories in CRC organoids. **A** UMAPs of organoid single cell transcriptomes, color coded by treatment conditions, as indicated. **B** Dendrograms of transcriptome similarities across treatment conditions, per organoid line. Height of dendrogram represents is obtained by hierarchical clustering on the overlap of KNN neighborhoods across conditions. (for details, see Methods). **C-H** Gene expression and RNA velocity analysis of P013T organoids under DMSO and EGFR inhibitor conditions. **C** UMAPs of the two conditions, color coded by condition **D** Developmental trajectories of organoid cells in the DMSO and EGFR treatment condition, as determined by RNA velocity. Grey arrow indicates endpoint of DMSO condition. Red arrows indicate the two endpoints of developmental trajectories after EGFR inhibition. DMSO cells are color coded by latent time. Latent time cannot be defined for EGFR inhibited cells showing divergent trajectories **E**, **F**, **H** UMAPs of the two conditions, color coded by cell cycle phase, activities of selected gene expression signatures, or gene-specific expression moments, respectively. **G** Phase plot and latent time RNA moments for *AXIN2* **I** Model of prototypical CRC cell trajectories.

We next assessed developmental trajectories, by computing dynamical cell velocity and latent time from scSLAM data. Cells of five out of six organoid lines developed along a MAPK gradient, and one line, OT227, did not show any preferential direction of cell development as assessed by RNA velocity (Fig. EV4A). The LGR5-ISC Wnt-driven stem cell signature marked the start of cell trajectories only in the BRAF-mutant lines. Except for OT227, developmental trajectories ended with increased goblet cell-like gene expression.

We next analyzed developmental trajectories in P013T organoids, comparing control and the clinically relevant anti-EGFR treatment conditions (Fig. 6C, D). Comparison of dynamical cell velocities indicated that the formerly uniform developmental trajectory along the MAPK gradient became divergent after EGFR inhibitor treatment, with one group of CRC cells developing towards an endpoint similar to unperturbed growth (Fig. 6D, red arrow 1), while the remaining cells formed opposite trajectories (Fig. 6D, red arrow 2). Cycling cells in G2/S were diminished across the complete UMAP space inhabited by EGFR inhibitor-treated cells (Fig 6E). The former developmental endpoint 1 was characterized by goblet cell differentiation, including the goblet cell marker TFF3 and apoptosis signatures (Fig 6F), while the latter endpoint 2 was characterized by high LGR5-ISC expression. Accordingly, endpoint 2 showed *de-novo* expression of the Wnt target *AXIN2* as indicated by the dynamics in the phase plot after EGFR inhibition (Fig. 6G), and also by the more restricted fields of expression (also termed moments) for the older non-labelled as compared to the newer labelled RNAs (Fig. 6H). These patterns are strong indications for the re-routing of cell development towards a state high in expression of Wnt-driven stem cell gene expression and agree with organoid culture showing slow outgrowth of resistant P013T colonies under long-term EGFR treatment (Fig. EV5).

We also observed characteristic changes in directed cell development after MAPK inhibition in most other CRC organoid lines (Fig. EV4B-D). RAS/RAF-wildtype P009T organoid cells likewise developed towards a state of increased LGR5-ISC-related gene expression after EGFR inhibition (Fig. EV4B). KRAS-mutant OT227 organoid cells, while displaying no clear trajectories in the absence of treatment, formed two streams of preferred trajectories after 48h of MEK inhibition, ending in regions of G1-arrested cells with active LGR5-ISC, differentiation and apoptosis transcriptome signatures (Fig. EV4C). In contrast, trajectories of OT302 organoid cells under MEK inhibition headed towards a region defined by goblet cell-like differentiation and apoptosis with decreased LGR5-ISC-related gene expression (Fig. EV4D). BRAF-mutant B2040 organoid cells developed towards a differentiation/apoptosis high trajectory endpoint under control conditions, but towards a LGR5-ISC-high state under combined EGFR/BRAF inhibition (Fig. EV10E). Taken together, in most lines examined, we observed separation of transcriptomes in the UMAP between control and effective treatment conditions, as a result of gene expression changes that can be interpreted as a re-routing of cell developmental trajectories towards endpoints defined either by differentiation followed by apoptosis, or by the induction of LGR5-ISC related gene expression characteristic for intestinal stem cells.

## Discussion

Here, we provide a comprehensive analysis of patient-overarching states of CRC cells forming developmental trajectories (Fig. 6 I). We define six clusters comprising cells resembling normal stem/TA cells, goblet cell-like tumor cells, and clusters TC1-4 that were largely confined to tumor tissue. The TC1-4 clusters were defined by differential activities of oncogenic pathways, including MAPK. We find that cell hierarchies in CRC are organized along developmental trajectories following MAPK gradients. In combination, the analyses imply that CRC cells develop along transient developmental states rather than forming fixed cell populations in organoids and probably also in patients. As cell state prevalence differed between patients, we suggest that CRC cell trajectories are guided and constrained by individual cancer genome characteristics such as patterns of oncogenic driver mutations. Our experiments in organoids indicate that therapies targeting the MAPK pathway reduce proliferation, but also redirect developmental trajectories of CRC cells towards goblet-cell differentiation-like apoptotic or Wnt-driven stem-cell like endpoints that could be associated with therapy-sensitivity or resistance, respectively. In summary, our analysis provides a single cell-based framework for cell plasticity during cancer development and targeted therapy.

Previous single cell studies defined features of the CRC immune microenvironment (James *et al*, 2020; Zhang *et al*, 2020; Lee *et al*, 2020); however, no consensus exists on patient-overarching features defining cells of the epithelial CRC compartment. Here, we define key states of CRC cells that are products of differential oncogene-associated signals resulting in distinctive transcriptomes. Our analyses suggest that CRC cells with lower levels of MAPK activity can resemble normal stem cells characterized by high expression of Wnt-driven LGR5-ISC signature genes (van de Wetering *et al*, 2002; Merlos-Suárez *et al*, 2011). However, individual stem cell markers such as *OLFM4* or *CD44* were expressed even stronger in the tumor-specific TC1-4 clusters. Based on these transcriptional patterns, it appears that no unique stem cell signature exists in CRC and that Wnt, YAP and MAPK activities together can maintain different cell states that may act as functional equivalents of stem cells that can sustain tissue growth. A recent study identified ribosomal transcription and protein translation as a key determinator of CRC stemness (Morral *et al*, 2020); this model agrees with our data, as ribosomal genes were overrepresented among the top-scoring genes determining the MAPK-driven trajectories described here (see Supplementary Fig. 5B for an example).

MAPK is a key pathway for targeted therapy, as many CRC patients profit from anti-EGFR or anti-EGFR/anti-BRAF therapy (Karapetis *et al*, 2008; Amado *et al*, 2008; Kopetz *et al*, 2019). By and large, outcomes of our experimental inhibition of MAPK in organoids agreed with known relationships between predictive mutations and therapy sensitivity, as EGFR inhibition was only effective in MAPK-wildtype organoids, while a combination of BRAF and EGFR inhibitors – but not each inhibitor alone – had profound effects on development of BRAF^V600E^-mutant CRC organoids. In addition, we show that graded MAPK-driven gene expression drives differentiation trajectories, as inferred from RNA velocity after MAPK inhibition in CRC organoids, extending our previous finding of graded ERK activity in CRC organoids (Brandt *et al*, 2019). In combination, these findings suggest that intrinsic resistance to anti-MAPK therapies may rely on re-routing of developmental trajectories of CRC cells. Indeed, the ability to reverse developmental trajectories in the intestinal epithelium has been found before (Schwitalla *et al*, 2013; Buczacki *et al*, 2013; Jadhav *et al*, 2017). Our study therefore adds new aspects to current models of anti-MAPK therapy resistance defined by cell plasticity (Woolston *et al*, 2019; Misale *et al*, 2014).

Importantly our study suggests that multiple signaling pathways form a therapy-relevant interconnected network of oncogenic signaling pathways in CRC. For instance, we find that YAP and MAPK levels are positively correlated on the protein activity and the transcriptional response levels. YAP maintains regenerative responses and is a key driver of CRC and other cancers (Zanconato *et al*, 2016). In contrast, MAPK and Wnt signaling were negatively correlated. We and others previously showed loss of Wnt-driven intestinal stem cells by high MAPK levels provided by oncogenic BRAF (Riemer *et al*, 2015; Tong *et al*, 2017). Here, we find that therapeutic inactivation of MAPK can result in Wnt and LGR5-ISC signature reactivation, in agreement with a previous study (Zhan *et al*, 2019). As both pathways, Wnt and MAPK, are generally activated by oncogenic mutations in CRC, cross-inhibition between the pathways would mean that therapeutical suppression of one pathway results in oncogene-driven activation of the other, possibly explaining why many therapeutic approaches to block MAPK proved insufficient in the clinic. It will be an important goal for future studies to identify combinations of actionable signals that can be exploited for therapies resulting in uniform commitment of CRC cells to differentiation-related and apoptotic endpoints instead of re-routing subsets of cancer cells towards stem cell-like states.

Our study also identified further cancer traits in CRC cell clusters with relevance to therapy. For instance, TC1 cells were defined by high levels of replication stress, which can be functionally associated with high MAPK activity (Sheu *et al*, 2012; Klotz-Noack *et al*, 2020).Tumors with high TC1 cell content were strongly positive for PARP, an important therapeutic target (Sun *et al*, 2020). It is of note that the CMS subtyping system developed for bulk tissue CRC transcriptomes (Guinney *et al*, 2015) could not distinguish the CRC cell types that we identified here on the single cell level, as most epithelial cancer cells were assigned to CMS1 or CMS2 with the exception of goblet-like cells that can adopt CMS3 (Supplementary Fig. 7).

In addition to patient-overarching CRC cell traits, we also observed patient-specific gene expression differences. Our integrated analysis of single cell transcriptomes and copy number gains and losses indicated that patient-specific gene expression patterns were significantly associated with copy number gains that we inferred from transcriptomes and validated using exome sequencing of three patients. On another level, we observed patient-specific gene expression patterns that translated into regional patterns of proteins, as evidenced by the staining for MMP7 which was confined to the invasive front of some tumors. Such patterns of gene activity and protein translation indicate that tumor cell development is highly plastic and partly regulated by immune and stromal cells in the microenvironment. Thus, we expect that the intrinsic developmental paths of CRC cells that we observe in organoid cultures are modulated by extrinsic cues from the microenvironment *in vivo*. Future studies using co-cultures of different tumor-associated cell types could disentangle key paracrine relationships in cancer.

We analyzed here primary cancer epithelium and organoid transcriptomes. Novel single cell approaches, taking into account the diversity of the tumor microenvironment in patient cohorts stratified by treatment, complex cell culture models, the extension of single-cell analyses to multi-omics, and the preservation of spatial information at a cellular level promise to identify cellular heterogeneity and genetic diversity of cancer at even greater detail in the future. The combination of such approaches has the potential to improve the molecular understanding of cancer and therapy prediction for patients (Rajewsky *et al*, 2020). Our work defining developmental trajectories of CRC contributes to this goal.

## Materials and Methods

### Collection and single-cell RNA sequencing of clinical specimens

Fresh normal colon and colorectal cancer tissues were acquired during the intraoperative pathologist’s examination at Charité University Hospital Berlin. Tissues (approx. 0.1-0.4g) were minced using scalpels, processed using the Miltenyi Human Tumor Dissociation Kit (Miltenyi, #130-095-929) and a Miltenyi gentleMACS Tissue Dissociator (Miltenyi, #130-096-427), using program 37C_h_TDK_1 for 30-45min. For three tumors, we also used digestion with the cold active protease from Bacillus licheniformis (Sigma P5380) at approx. 6°C for 45min. with frequent agitation, following a published protocol (Adam *et al*, 2017) (Supplementary Fig 8). Cell suspensions were filtered using 100μm filters, pelleted by centrifugation, treated with 1ml ACK erythrocyte lysis buffer, washed and resuspended in ice-cold PBS, and filtered using 20μm filters. Debris was removed using the Debris Removal Solution (Miltenyi #130-109-398). Cell suspensions were analyzed for cell viability >75% using LIVE/DEAD Fixable Dead Cell Stain Kit (488nm; Thermo Fisher) and a BD Accuri cytometer. 10.000 single cells were used for single-cell library production, using the Chromium Single Cell 3’Reagent Kits v3 and the Chromium Controller (10x Genomics). Libraries were sequenced on a HiSeq 4000 Sequencer (Illumina) at 200-400mio. reads per library to a mean library saturation of approx. 50%. This resulted in 35.000 to 120.000 reads per cell.

### DNA Sequencing

For panel sequencing, DNA was extracted from FFPE tumor tissue using the Maxwell RSC DNA FFPE Kit (Promega) or the GeneRead DNA FFPE kit (Qiagen) and sequenced using a CRC panel (Mamlouk *et al*, 2017), and/or the Ion AmpliSeq Cancer Hotspot Panel (CHP) v2 and an IonTorrent sequencer (ThermoFisher). Variant calling was performed using Sequence Pilot (Version 4.4.0, JSI Medical Systems) or SoFIA (Mamlouk *et al*, 2017). For exome sequencing, DNA was isolated from fresh frozen tumor tissue using the DNeasy Blood and Tissue Kit (Qiagen). Exomes were sequenced using the AllExon Human SureSelect v7 Kit (Agilent).

### Histology and immunostaining

3-5 μm tissue sections of formalin-fixed and paraffin-embedded (FFPE) tissue were used. Immunostainings were performed on the BenchMark XT immunostainer (Ventana Medical Systems), using CC1 mild buffer or Ultra CC1 buffer (Ventana Medical Systems) for 30 min at 100°C, and using antibodies rabbit anti-TFF3 (1:250, Abcam, ab108599), mouse anti-FABP1 (1:1000, Abcam, ab7366), rabbit anti-OLFM4 (1:100, Atlas Antibodies, HPA077718), mouse anti-EPCAM (1:100, ThermoScientific, MS-144-P1), rabbit anti-Ki67 (1:400, Abcam, ab16667), mouse anti-Ki67 (1:50, Dako, M7240), rabbit anti-LYZ (1:1500, Abcam, ab108508), rabbit anti-EREG (1:50, Thermo Fischer Scientific, PA5-24727), anti-PARP1, mouse anti-MUC2 (1:50, Leica, NCL-MUC-2), mouse anti-CK17 (1:10, Dako, M7046), mouse anti-MMP7 (1:100, ThermoFisherScientific, MA5-14215). Images were taken using AxioVert.A1 (Zeiss) or CQ1 (Yokogawa) microscopes or scanned using the Pannoramic SCAN 150 scanner (3DHISTECH).

### Organoid culture and metabolic labelling

Tumor cells were washed in Advanced DMEM/F12 medium (Gibco), embedded in Matrigel, and cultured in 24-well plates, as published (Sato *et al*, 2011). Wnt3 and R-Spondin3 were prepared as conditioned media (Sato *et al*, 2011). For single-cell sequencing, organoids were dissociated completely using TrypLE and DNAseI, and filtered via a 20μm filter. For single cell SLAM-seq, NCO, P009T and P013T replicate cultures were cultured in media with and without Wnt/R-Spondin, and P009T, P013T, OT227, OT302, B2040, C2019 organoids were cultured in standard media (Sato *et al*, 2011; Schütte *et al*, 2017) with DMSO, or were treated for 48 with 100nM AZD8931 (Sapatinib), 100nM LGX818 (Encorafenib) and/or 8μM AZD6244 (Selumetinib). Organoids were metabolically labelled in culture using 200μM 4-thio-uridine for 2 h (Herzog *et al*, 2017), harvested, disaggregated to single cells by TrypLE, and fixed in fixation buffer (80% methanol/20% DPBS) at ≥ −20° C. Samples were warmed to room temperature and incubated with 10mM iodoacetamide. Alkylation was carried out overnight, in the dark, with gentle rotation, followed by two washes with cold fixation buffer. Single cell suspensions were rehydrated and incubated 10 minutes at room temperature in 100mM DTT. Samples were resuspended in fixation buffer and conserved at −80 °C.

### Mass Cytometry (CyTOF)

For CyTOF analysis, we used a panel of antibodies described in Brandt *et al*, 2019, and measurements were performed essentially as in described in the same publication. In short, organoids were harvested in PBS and digested to a single cell solution in 1:1 Accutase (Biolegend) and TrypLE (Gibco) with addition of 100 U/ml Universal Nuclease (Thermo Scientific) at 37°C. Cells were counted and a maximum of 500.000 cells were stained with 5 μM Cell-ID Cisplatin (Fluidigm) in PBS for 5 min at 37°C, washed in PBS, resuspended in medium and incubated for 30 min at 37°C, resuspended in BSA/PBS solution, mixed 1:1.4 with Proteomics Stabilizer (Smart Tube Inc.) and frozen at −80°C for storage.

For analysis, cells were thawed, mixed with Maxpar Cell Staining Buffer (CSB, Fluidigm), labelled using the Cell-ID 20-Plex Pd Barcoding Kit, and washed again in CSB, then in Barcode Perm Buffer (Fluidigm). After barcoding, cells were pooled and stained with a surface antibody cocktail, as described previously (Brandt *et al*, 2019). Data was acquired on a Helios CyTOF system. Mass cytometry data was normalized using the Helios software and bead-related events were removed. Doublets were excluded by filtering for DNA content (^191^Ir and ^193^Ir) vs. event length, and apoptotic debris removed by a filter in the platin channel (^195^Pt). De-convolution of the barcoded sample was performed using the CATALYST R package version 1.5.3 (Chevrier *et al*, 2018).

### Primary tissue single-cell RNA-seq data analysis

UMIs were quantified using cellranger 3.0.2 with reference transcriptome GRCh38. Spliced, unspliced and ambiguous UMIs were quantified with velocyto (La Manno *et al*, 2018) (mode: run10x, default parameters). For parameters and initial quality controls, see Supplementary Fig. 2. Epithelium, stromal, and immune cells were identified by scoring cell type markers across Louvain clusters for each sample (resolution = 1). Cell type markers used to score epithelium, stromal, and immune cells were adapted from Similie et al. (Smillie *et al*, 2019) and are listed in Supplementary table 3. Sample-wise quality control assessments and subsettings into main cell types are documented at sys-bio.net/sccrc/. Normalized subsets were merged for each main cell type of normal and tumor samples without further batch correction.

SNN graph, Louvain clusters and UMAP embeddings were recomputed for each subset based on top ten components. Louvain cluster-specific marker genes of merged normal and tumor samples were used to identify sub cell types among epithelial, stromal and immune subsets. Here, marker genes were determined with Seurat (Stuart *et al*, 2019)(wilcox text) at a minimum log fold change threshold of 0.25. Gene expression sets were taken from the hallmark signature collection of the Broad institute (Liberzon *et al*, 2015), unless otherwise referenced in the main text, and were scored as implemented in the progeny R package and Seurat v3, respectively.

For copy-number assessment, InferCNV v1.3.3 was used with default parameters. Copy number aberrant clones were cut at k = 2 in inferCNV dendrograms. Clone-wise SCNA scores were computed by calculating the average standard deviation in inferCNV expression of all cells and divided by the average standard deviation in inferCNV expression of all normal samples taken together. Clones with a SCNA score greater than the highest observed score for normal samples were considered copy number aberrant.

### Organoid single-cell RNA-seq data analysis

Single cell SLAM sequencing data was pre-processed using cellranger v4.0, and labeled and unlabeled reads were counted using the alignments (as BAM files), using a custom pipeline utilizing Snakemake (Köster & Rahmann, 2012), SeqAn (Döring *et al*, 2008), R. For each read, the numbers of T nucleotides and T-to-C conversions was counted, leaving out positions with common SNPs (using the dbSNP build 151 as available as track from UCSC genome browser). For each molecule as identified by cell barcode and UMI, positions with discordant nucleotides were excluded. Subsequently, molecules were counted as nascent RNA if they contained a T-to-C conversion, and old RNA otherwise.

scRNAseq and scSLAM-seq data for organoids was analyzed using scanpy (Wolf *et al*, 2018)and scvelo (Bergen *et al*, 2020). For diffusion map analysis and RNA velocity, cells were first filtered by the number of genes (between 2000 and 5000) and the percent mitochondrial reads (between 0.075 and 0.2) and normalized, using scvelo standard settings. Cell cycle was scored according to the scanpy standards. Cell cycle was scored based on a list of cell cycle associated genes (Kowalczyk *et al*, 2015). Resulting S_score, G2M_score and UMI counts per cell were regressed out. The diffusion map was calculated on the top 10 components and using a neighborhood graph with 50 neighbors and calculated on all genes. UMAP embeddings were computed based on a PCA on only the 2000 most highly variable genes obtained with scanpy. The similarity measure between conditions in Figure 6B was defined by the average fraction of shared neighbors in a nearest neighbor graph over all cells in the dataset, then performing hierarchical clustering on the resulting similarity matrix.

Moments were calculated on 30 principal components and 30 neighbors separately per condition in order to avoid smoothing effects between different conditions within the same dataset. Velocity was calculated using the dynamical model from scvelo on 2000 most highly variable genes according to scanpy, which were then filtered for at least 20 shared counts in both SLAM layers. Furthermore, for most of the captured genes in the first perturbation experiment we observed either induced or repressed gene dynamics only. Such one-sided dynamics proved ambiguous to fit kinetics to, since at least two local maxima of model likelihood are possible. In order to resolve this ambiguity, we modified the dynamical model from scvelo with an additional regularization term. This term introduces prior information to the model by penalizing the number of cells that are assigned a higher latent time than a given set root priors. Before fitting we inferred the root cells in a data-driven manner as those ten cells with the stem cell signature. An additional initialization of each gene model with restricted parameters ensured that both local minima were actually captured. When fitting the second perturbation experiment datasets we did not observe any benefit when imposing a prior, hence continued without one.

## Ethics permission

All patients were aware of the planned research and agreed to the use of tissue. Research was approved by vote EA4/164/19 of the ethic’s commission of Charité - Universitätsmedizin Berlin.

## Data availability

Scripts for processing of patient tissue scRNA sequencing data are available from https://github.com/molsysbio/sccrc. Scripts for processing of organoid RNA velocity data are available from https://github.com/molsysbio/sccrc_slamvelocity. Raw data is stored in GEO under the accession numbers GSE166555 (patient data) und GSE166556 (organoid data). Processed count data is available from https://sys-bio.net/sccrc.

## Acknowledgements

The work was in part funded by Berlin Institute of Health (PB, NB, CS, DH and MM), German Cancer Consortium DKTK (NB, MM), Deutsche Forschungsgemeinschaft (MM, grant MO2783/5), DFG research training group (SP and TTW, grant GRK2424/1), BMBF (NB), and the Helmholtz Association (RFS).

## Author contributions

PB, FU, SP, MM, AS, MLu, AT, TS, RA, YR, SM, KKN conducted and analyzed experiments; FU, SP, NB, BO, EB, TTW performed bioinformatic analyses; MM, NB, PB, CS, DB, BS, DH, MLa, RFS conceived, designed, interpreted experiments and/or supervised parts of the study; PB, MM, DH, CK, AT contributed to clinical sample acquisition and preparation; MM wrote the manuscript; all authors provided critical feedback and helped shaping the research, analysis, and manuscript.

## The Paper Explained

### PROBLEM

Cancer cells, like normal cells in our bodies, are thought to develop along preferred trajectories. It is unknown which signals control development of colorectal cancer cells, and how cancer therapies interfere with such signals.

### RESULTS

We find that colorectal cancer cells develop along trajectories defined by activity of a signaling pathway termed MAPK, which is also a clinically important target for therapy. We find that experimental therapies inactivating MAPK disturb development of colorectal cancer cells in culture, resulting in cancer cells heading towards an unwanted stem cell-like state. This cellular behavior might explain cases of therapy resistance observed in patients.

### IMPACT

Knowledge of developmental routes of cancer cells is key for future improvement of targeted therapy to avoid resistance.

## Extended View Figures

**Figure EV1:**
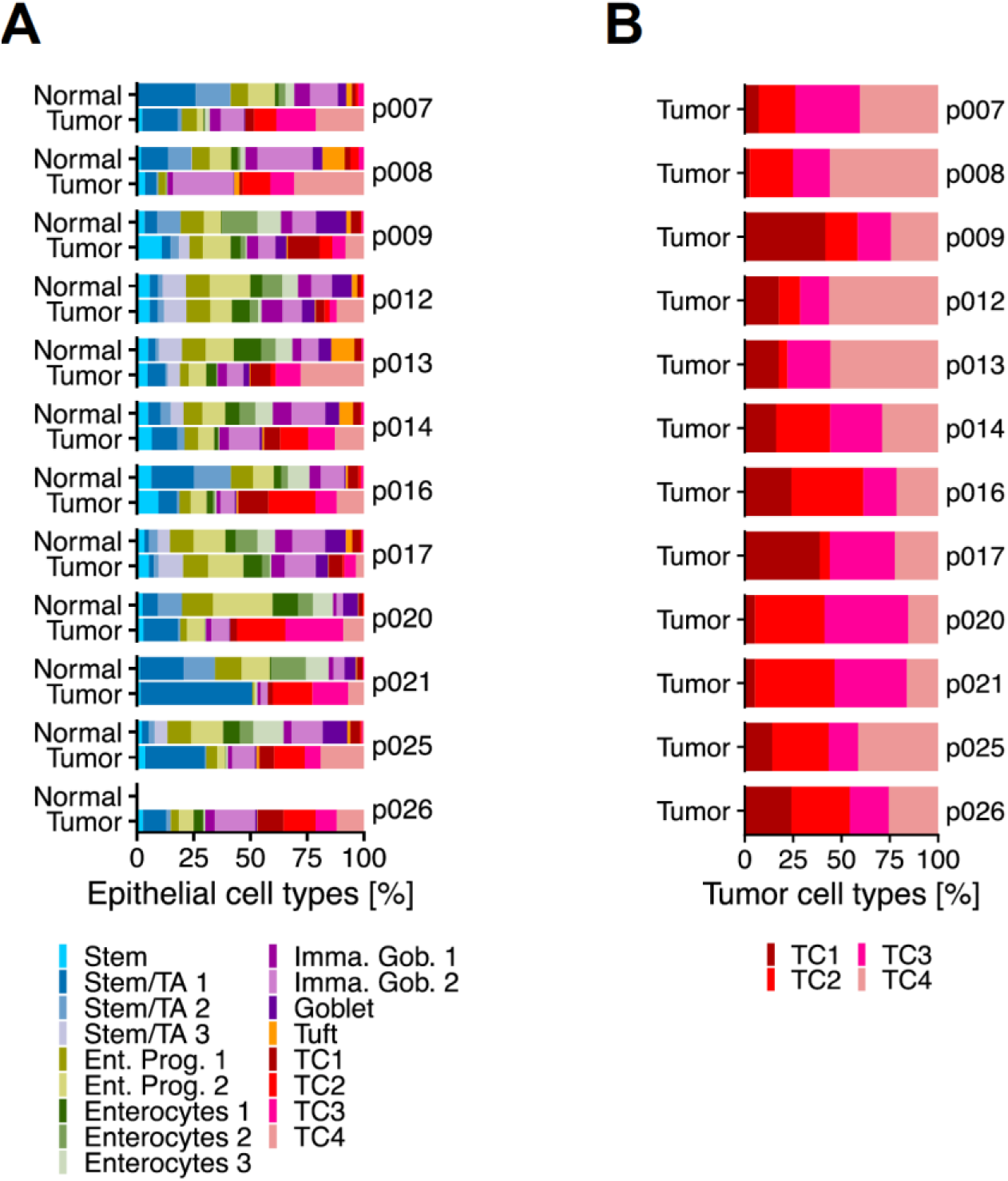
Cell type census, per patient. **A** Relative prevalence of cells in the epithelial compartment, per patient **B** Relative prevalence of TC1-4 cluster cells, per patient.

**Figure EV2:**
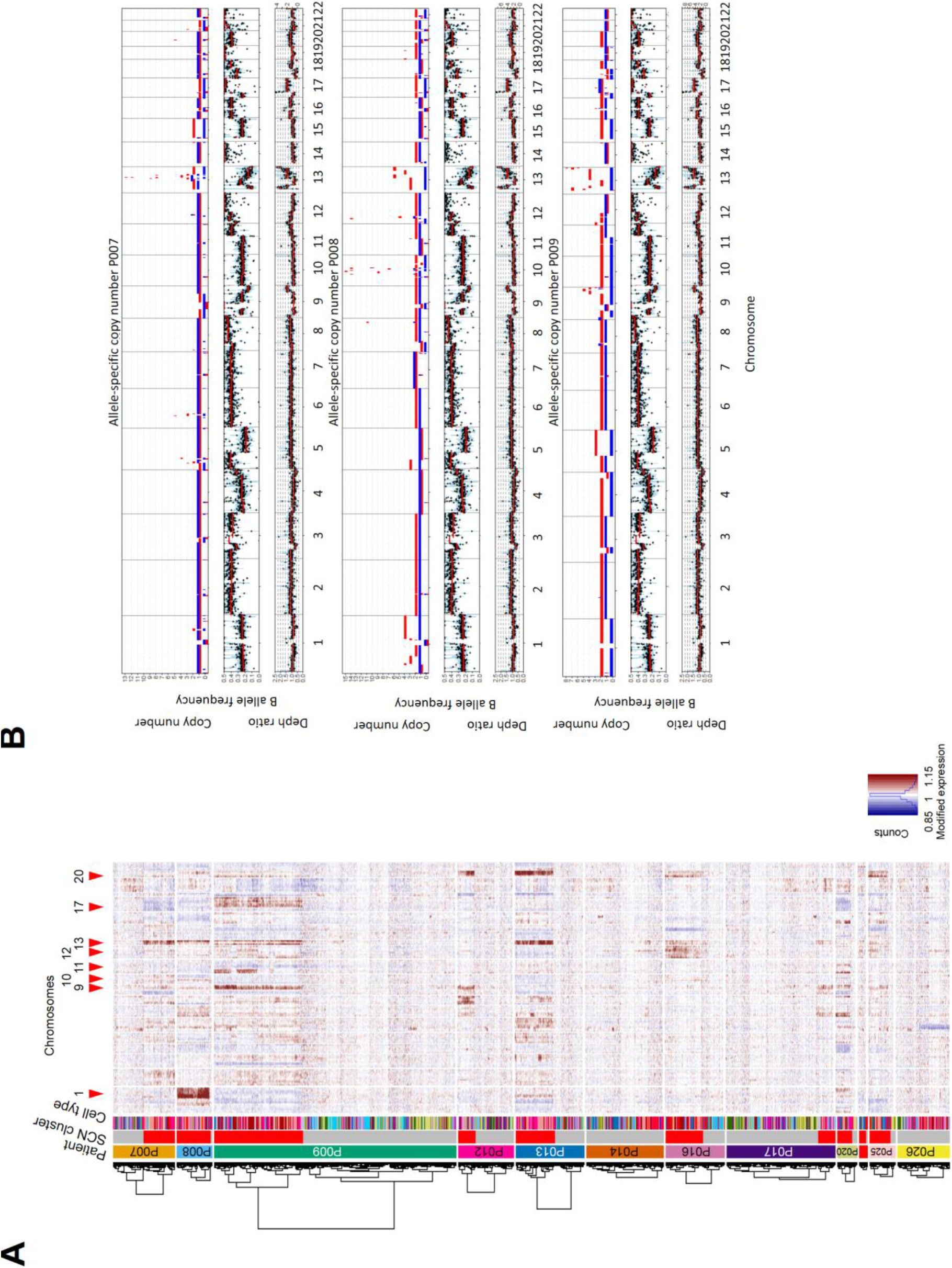
Assessment of copy-number variations. **A** Copy-number calling from single cell transcriptomes, using InferCNV. **B** Copy-number calling from exome sequences. To validate SCNAs from single cell transcriptome data, we performed allele-specific SCNA calling for patients P007, P008 and P009 from bulk whole exome data. Germline variants were discovered *de novo* and read counts were accumulated for each allele at heterozygous germline variants using bcftools v1.9 multi-allelic caller. Discovered variants and read counts were passed to Sequenza 3.0 for segmentation and calling of allele specific SCNAs. Top tracks: copy number segments of major and minor alleles. Middle tracks: B (minor) allele frequency. Bottom tracks: depth ratio of matched tumor versus normal samples with corresponding second axis of inferred copy number. P007 showed multiple LOH events on chromosomes 1, 3, 5, 6, 8, 9, 13, 15, 17 and 21 as well as a focal amplification on chromosome 13. P008 demonstrated a similar SCNA pattern with LOH events on chromosomes 8, 9, 10, 12, 13, 17 and 22 as well as multiple focal amplifications on chromosomes 8, 10, 12 and 13. P009 was characterized by widespread copy-number neutral LOH in about 50% of the genome. Focal amplifications were found on chromosomes 9 and 13.

**Figure EV3:**
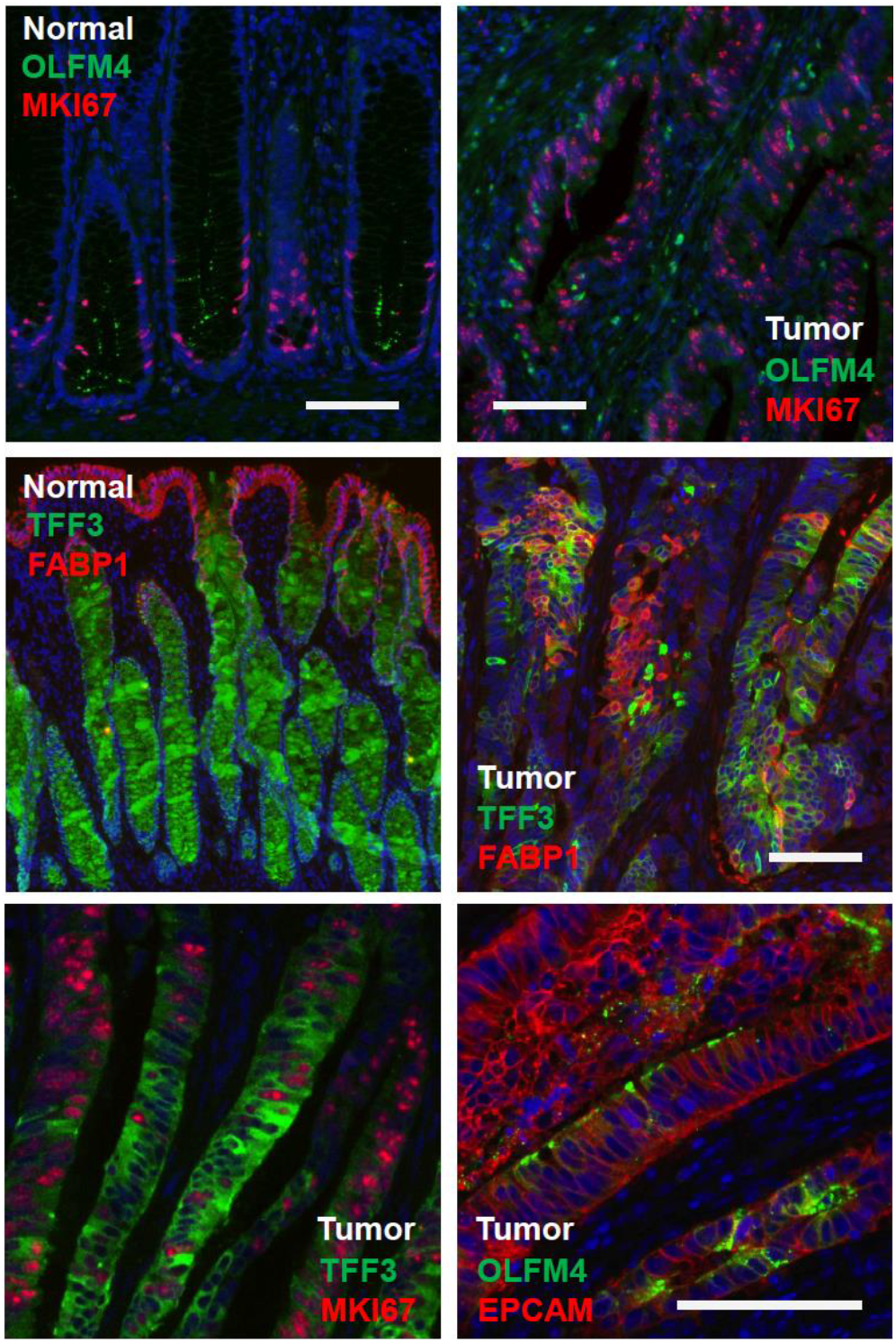
Assessment of CRC cell heterogeneity by immunofluorescence. Immunofluorescence analysis for OLFM4, MKI67, FABP1, and TFF3 in normal and tumor tissue. All sections are from patient P009, except the EPCAM/OLFM4 co-staining that was done on tumor tissue of P016. Scale bars indicate 100μm. For marker expression in UMAP space, see Supplementary Fig. 3, above.

**Figure EV4:**
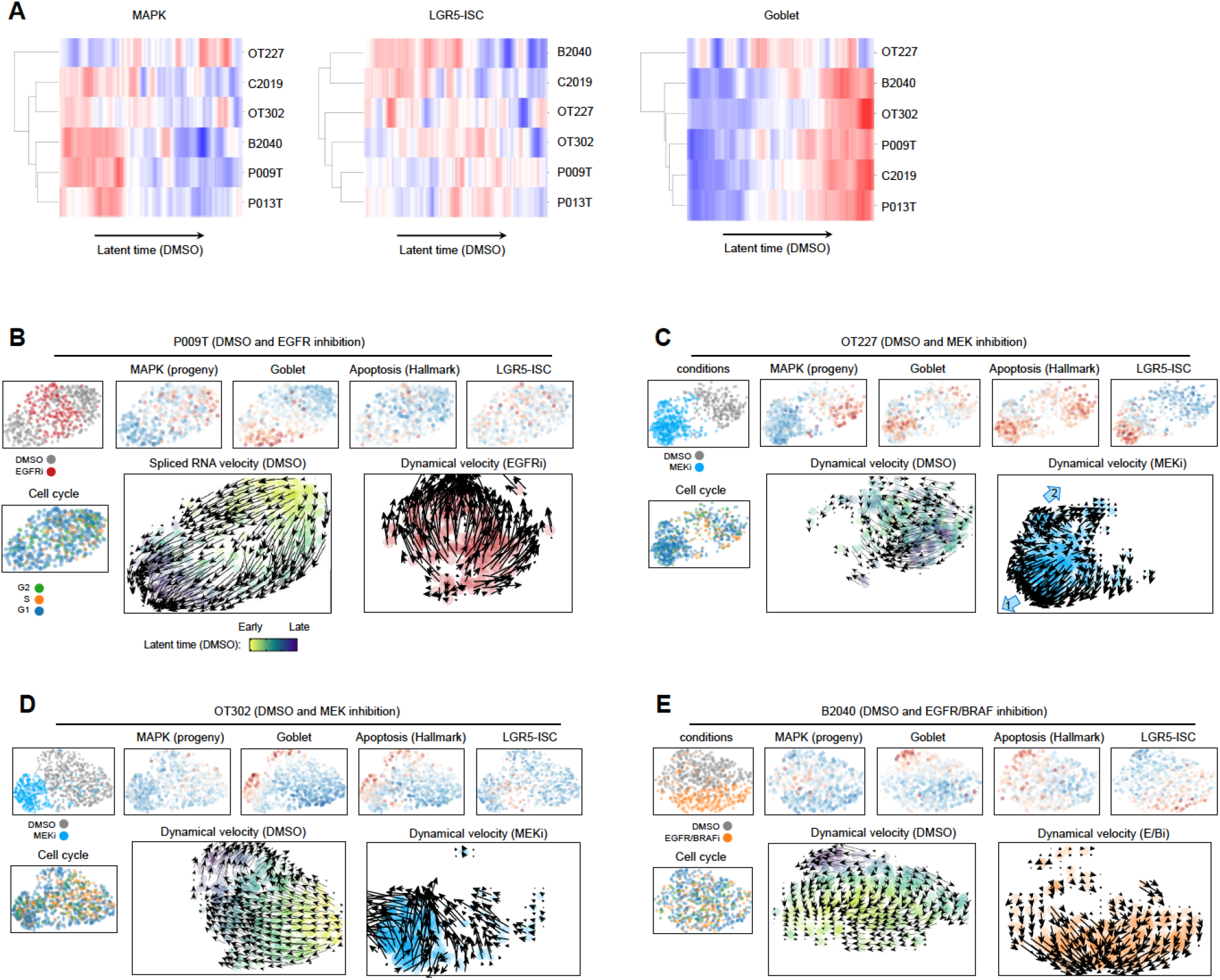
Analysis of latent time and RNA velocity in CRC organoids. **A** Strength of MAPK, LGR5-ISC and Goblet-related signature gene expression along latent time, per line in control conditions (that is, without anti-MAPK treatment). No uniform trajectories could be called for OT227, resulting in random latent time. **B** Gene expression and RNA velocity analysis of P009T organoids under DMSO and EGFR inhibitor conditions. For this line, calling of RNA velocities ran in opposite directions for dynamical RNA velocity based on SLAM information, and for classical steady-state velocity by splicing. Only the latter velocity was in agreement with latent time. Based on analysis of phase plots, we propose that latent time and splicing-based RNA velocity are correct and represent the underlying biology, while SLAM velocity models may be confounded by genes displaying a OFF dynamic in disagreement with latent time. **C-D** Gene expression and RNA velocity analysis of OT227 and OT302 organoids under DMSO and MEK inhibitor conditions. **E** Gene expression and RNA velocity analysis of B2040 organoids under DMSO and combined EGFR/BRAF inhibitor conditions.

**Figure EV5:**
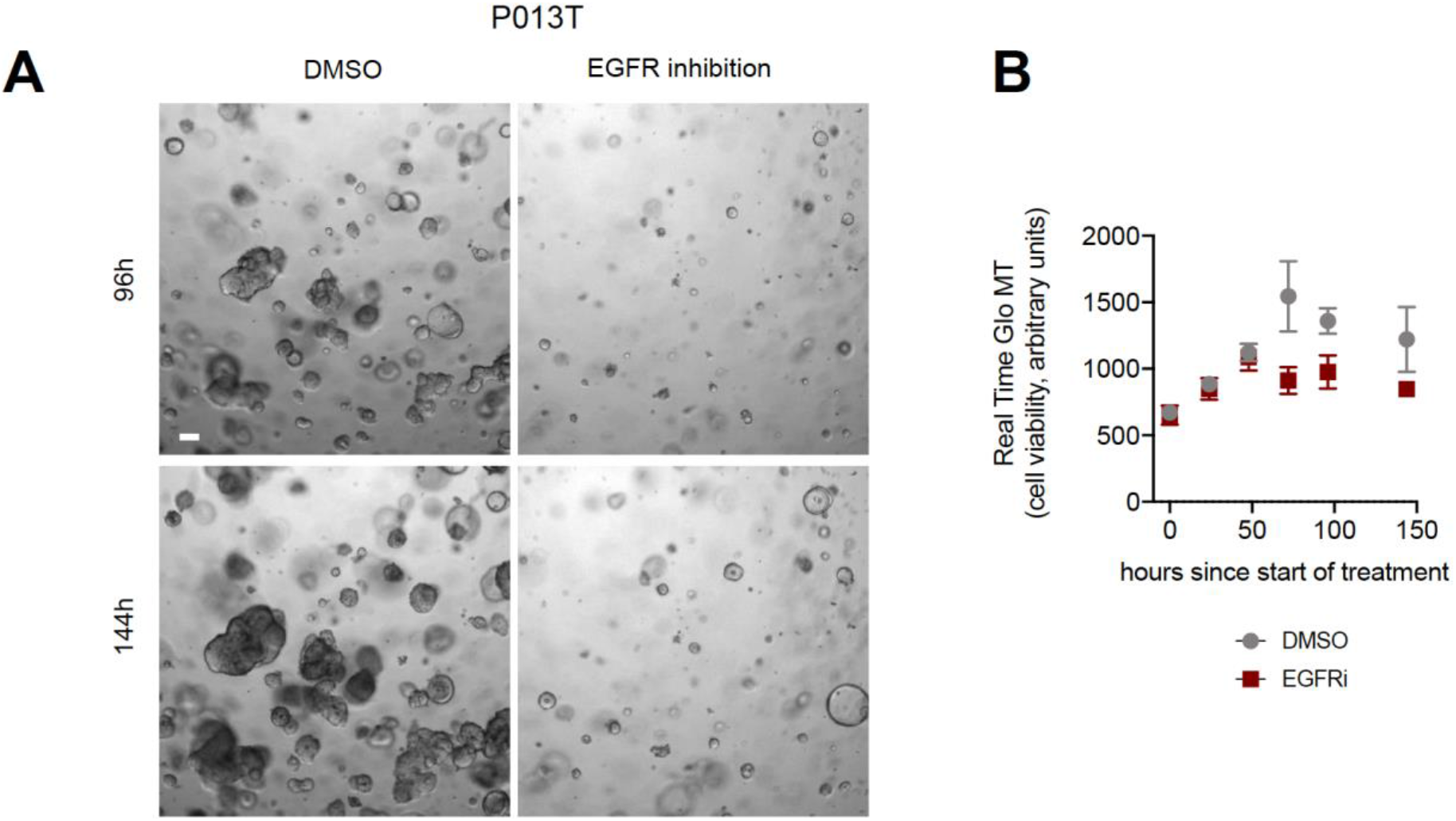
Long-term organoid culture of P013T reveals outgrowth of EGFR-resistant cells. **A** Phenotypes of DMSO control and EGFR-treated P013T cultures after 96 and 144h, as indicated. Organoids growing under EGFR inhibition show a distinct spheroidal phenotype. **B** Metabolic cell viability assays of triplicate DMSO control and EGFR inhibitor treated P013T cultures.

## References

Adam M, Potter AS & Potter SS (2017) Psychrophilic proteases dramatically reduce single-cell RNA-seq artifacts: A molecular atlas of kidney development. Dev 144: 3625–3632

Amado RG, Wolf M, Peeters M, Van Cutsem E, Siena S, Freeman DJ, Juan T, Sikorski R, Suggs S, Radinsky R, et al (2008) Wild-type KRAS is required for panitumumab efficacy in patients with metastatic colorectal cancer. J Clin Oncol 26: 1626–1634

Bergen V, Lange M, Peidli S, Wolf FA & Theis FJ (2020) Generalizing RNA velocity to transient cell states through dynamical modeling. Nat Biotechnol 38: 1408–1414

Brabletz T, Jung A, Dag S, Hlubek F & Kirchner T (1999) β-catenin regulates the expression of the matrix metalloproteinase-7 in human colorectal cancer. Am J Pathol 155: 1033–1038

Brandt R, Sell T, Lüthen M, Uhlitz F, Klinger B, Riemer P, Giesecke-Thiel C, Schulze S, El-Shimy IA, Kunkel D, et al (2019) Cell type-dependent differential activation of ERK by oncogenic KRAS in colon cancer and intestinal epithelium. Nat Commun 10: 2919

Buczacki SJA, Zecchini HI, Nicholson AM, Russell R, Vermeulen L, Kemp R & Winton DJ (2013) Intestinal label-retaining cells are secretory precursors expressing Lgr5. Nature 495: 65–69

Chevrier S, Crowell HL, Zanotelli VRT, Engler S, Robinson MD & Bodenmiller B (2018) Compensation of Signal Spillover in Suspension and Imaging Mass Cytometry. Cell Syst 6: 612–620

Corcoran RB, Andre T, Atreya CE, Schellens JHM, Yoshino T, Bendell JC, Hollebecque A, McRee AJ, Siena S, Middleton G, et al (2018) Research article combined BRAF, EGFR, and MEK inhibition in patients with BRAF V600E -mutant colorectal cancer. Cancer Discov 8: 428–443

Van Cutsem E, Köhne C-H, Hitre E, Zaluski J, Chang Chien C-R, Makhson A, D’Haens G, Pintér T, Lim R, Bodoky G, et al (2009) Cetuximab and chemotherapy as initial treatment for metastatic colorectal cancer. N Engl J Med 360: 1408–1417

Döring A, Weese D, Rausch T & Reinert K (2008) SeqAn An efficient, generic C++ library for sequence analysis. BMC Bioinformatics 9: 11

Fearon ER (2011) Molecular genetics of colorectal cancer. Annu Rev Pathol 6: 479–507

Francescangeli F, Contavalli P, De Angelis ML, Careccia S, Signore M, Haas TL, Salaris F, Baiocchi M, Boe A, Giuliani A, et al (2020) A pre-existing population of ZEB2+ quiescent cells with stemness and mesenchymal features dictate chemoresistance in colorectal cancer. J Exp Clin Cancer Res 39

Fritsche-Guenther R, Witzel F, Sieber A, Herr R, Schmidt N, Braun S, Brummer T, Sers C & Blüthgen N (2011) Strong negative feedback from Erk to Raf confers robustness to MAPK signalling. Mol Syst Biol 7: 489

Goto N, Fukuda A, Yamaga Y, Yoshikawa T, Maruno T, Maekawa H, Inamoto S, Kawada K, Sakai Y, Miyoshi H, et al (2019) Lineage tracing and targeting of IL17RB+ tuft cell-like human colorectal cancer stem cells. Proc Natl Acad Sci U S A 116: 12996–13005

Guinney J, Dienstmann R, Wang X, de Reyniès A, Schlicker A, Soneson C, Marisa L, Roepman P, Nyamundanda G, Angelino P, et al (2015) The consensus molecular subtypes of colorectal cancer. Nat Med 21: 1350–1356

Hanahan D & Weinberg RA (2011) Hallmarks of cancer: the next generation. Cell 144: 646–674

Hatzivassiliou G, Song K, Yen I, Brandhuber BJ, Anderson DJ, Alvarado R, Ludlam MJC, Stokoe D, Gloor SL, Vigers G, et al (2010) RAF inhibitors prime wild-type RAF to activate the MAPK pathway and enhance growth. Nature 464: 431–435

Van Der Heijden M, Miedema DM, Waclaw B, Veenstra VL, Lecca MC, Nijman LE, Van Dijk E, Van Neerven SM, Lodestijn SC, Lenos KJ, et al (2019) Spatiotemporal regulation of clonogenicity in colorectal cancer xenografts. Proc Natl Acad Sci U S A 116: 6140–6145

Herzog VA, Reichholf B, Neumann T, Rescheneder P, Bhat P, Burkard TR, Wlotzka W, Von Haeseler A, Zuber J & Ameres SL (2017) Thiol-linked alkylation of RNA to assess expression dynamics. Nat Methods 14: 1198–1204

Jadhav U, Saxena M, O’Neill NK, Saadatpour A, Yuan GC, Herbert Z, Murata K & Shivdasani RA (2017) Dynamic Reorganization of Chromatin Accessibility Signatures during Dedifferentiation of Secretory Precursors into Lgr5+ Intestinal Stem Cells. Cell Stem Cell 21: 65–77

James KR, Gomes T, Elmentaite R, Kumar N, Gulliver EL, King HW, Stares MD, Bareham BR, Ferdinand JR, Petrova VN, et al (2020) Distinct microbial and immune niches of the human colon. Nat Immunol 21: 343–353

Jürges C, Dölken L & Erhard F (2018) Dissecting newly transcribed and old RNA using GRAND-SLAM. Bioinformatics 34: i218–i226

Karapetis CS, Khambata-Ford S, Jonker DJ, O’Callaghan CJ, Tu D, Tebbutt NC, Simes RJ, Chalchal H, Shapiro JD, Robitaille S, et al (2008) K-ras Mutations and Benefit from Cetuximab in Advanced Colorectal Cancer. N Engl J Med 359: 1757–1765

Klotz-Noack K, Klinger B, Rivera M, Bublitz N, Uhlitz F, Riemer P, Lüthen M, Sell T, Kasack K, Gastl B, et al (2020) SFPQ Depletion Is Synthetically Lethal with BRAFV600E in Colorectal Cancer Cells. Cell Rep 32: 108184

Kopetz S, Grothey A, Yaeger R, Van Cutsem E, Desai J, Yoshino T, Wasan H, Ciardiello F, Loupakis F, Hong YS, et al (2019) Encorafenib, binimetinib, and cetuximab in BRAF V600E–mutated colorectal cancer. N Engl J Med 381: 1632–1643

Köster J & Rahmann S (2012) Snakemake-a scalable bioinformatics workflow engine. Bioinformatics 28: 2520–2522

Kowalczyk MS, Tirosh I, Heckl D, Rao TN, Dixit A, Haas BJ, Schneider RK, Wagers AJ, Ebert BL & Regev A (2015) Single-cell RNA-seq reveals changes in cell cycle and differentiation programs upon aging of hematopoietic stem cells. Genome Res 25: 1860–1872

Kreso A & Dick JE (2014) Evolution of the cancer stem cell model. Cell Stem Cell 14: 275–291

Lamprecht S, Schmidt EM, Blaj C, Hermeking H, Jung A, Kirchner T & Horst D (2017) Multicolor lineage tracing reveals clonal architecture and dynamics in colon cancer. Nat Commun 8: 1406

Lasry A, Zinger A & Ben-Neriah Y (2016) Inflammatory networks underlying colorectal cancer. Nat Immunol 17: 230–240

Lee HO, Hong Y, Etlioglu HE, Cho YB, Pomella V, Van den Bosch B, Vanhecke J, Verbandt S, Hong H, Min JW, et al (2020) Lineage-dependent gene expression programs influence the immune landscape of colorectal cancer. Nat Genet 52: 594–603

Liberzon A, Birger C, Thorvaldsdóttir H, Ghandi M, Mesirov JP & Tamayo P (2015) The Molecular Signatures Database Hallmark Gene Set Collection. Cell Syst 1: 417–425

Mamlouk S, Childs LH, Aust D, Heim D, Melching F, Oliveira C, Wolf T, Durek P, Schumacher D, Bläker H, et al (2017) DNA copy number changes define spatial patterns of heterogeneity in colorectal cancer. Nat Commun 8: 14093

La Manno G, Soldatov R, Zeisel A, Braun E, Hochgerner H, Petukhov V, Lidschreiber K, Kastriti ME, Lönnerberg P, Furlan A, et al (2018) RNA velocity of single cells. Nature 560: 494–498

McInnes L, Healy J, Saul N & Großberger L (2018) UMAP: Uniform Manifold Approximation and Projection. J Open Source Softw 3: 861

Merlos-Suárez A, Barriga FM, Jung P, Iglesias M, Céspedes MV, Rossell D, Sevillano M, Hernando-Momblona X, da Silva-Diz V, Muñoz P, et al (2011) The intestinal stem cell signature identifies colorectal cancer stem cells and predicts disease relapse. Cell Stem Cell 8: 511–524

Misale S, Di Nicolantonio F, Sartore-Bianchi A, Siena S & Bardelli A (2014) Resistance to Anti-EGFR therapy in colorectal cancer: From heterogeneity to convergent evolution. Cancer Discov 4: 1269–1280

Morral C, Stanisavljevic J, Hernando-Momblona X, Mereu E, Álvarez-Varela A, Cortina C, Stork D, Slebe F, Turon G, Whissell G, et al (2020) Zonation of Ribosomal DNA Transcription Defines a Stem Cell Hierarchy in Colorectal Cancer. Cell Stem Cell 26: 845–861

Muñoz J, Stange DE, Schepers AG, van de Wetering M, Koo B-K, Itzkovitz S, Volckmann R, Kung KS, Koster J, Radulescu S, et al (2012) The Lgr5 intestinal stem cell signature: robust expression of proposed quiescent ‘+4’ cell markers. EMBO J 31: 3079–3091

O’Brien CA, Pollett A, Gallinger S, Dick JE, O’Brien CA, Pollett A, Gallinger S & Dick JE (2007) A human colon cancer cell capable of initiating tumour growth in immunodeficient mice. Nature 445: 106–110

De Palma FDE, D’argenio V, Pol J, Kroemer G, Maiuri MC & Salvatore F (2019) The molecular hallmarks of the serrated pathway in colorectal cancer. Cancers (Basel) 11: 1017

Parikh K, Antanaviciute A, Fawkner-Corbett D, Jagielowicz M, Aulicino A, Lagerholm C, Davis S, Kinchen J, Chen HH, Alham NK, et al (2019) Colonic epithelial cell diversity in health and inflammatory bowel disease. Nature 567: 49–55

Rajewsky N, Almouzni G, Gorski SA, Aerts S, Amit I, Bertero MG, Bock C, Bredenoord AL, Cavalli G, Chiocca S, et al (2020) LifeTime and improving European healthcare through cell-based interceptive medicine. Nature 587: 377–386

Ricci-Vitiani L, Lombardi DG, Pilozzi E, Biffoni M, Todaro M, Peschle C & De Maria R (2007) Identification and expansion of human colon-cancer-initiating cells. Nature 445: 111–115

Riemer P, Sreekumar A, Reinke S, Rad R, Schäfer R, Sers C, Bläker H, Herrmann BG & Morkel M (2015) Transgenic expression of oncogenic BRAF induces loss of stem cells in the mouse intestine, which is antagonized by β-catenin activity. Oncogene 34: 3164–3175

Sato T, Stange DE, Ferrante M, Vries RGJ, van Es JH, van den Brink S, Van Houdt WJ, Pronk A, Van Gorp J, Siersema PD, et al (2011) Long-term expansion of epithelial organoids from human colon, adenoma, adenocarcinoma, and Barrett’s epithelium. Gastroenterology 141: 1762–1772

Schubert M, Klinger B, Klünemann M, Sieber A, Uhlitz F, Sauer S, Garnett MJ, Blüthgen N & Saez-Rodriguez J (2018) Perturbation-response genes reveal signaling footprints in cancer gene expression. Nat Commun 9: 20

Schütte M, Risch T, Abdavi-Azar N, Boehnke K, Schumacher D, Keil M, Yildiriman R, Jandrasits C, Borodina T, Amstislavskiy V, et al (2017) Molecular dissection of colorectal cancer in pre-clinical models identifies biomarkers predicting sensitivity to EGFR inhibitors. Nat Commun 8: 14262

Schwitalla S, Fingerle AA, Cammareri P, Nebelsiek T, Göktuna SI, Ziegler PK, Canli O, Heijmans J, Huels DJ, Moreaux G, et al (2013) Intestinal Tumorigenesis Initiated by Dedifferentiation and Acquisition of Stem-Cell-like Properties. Cell 152: 25–38

Serra D, Mayr U, Boni A, Lukonin I, Rempfler M, Challet Meylan L, Stadler MB, Strnad P, Papasaikas P, Vischi D, et al (2019) Self-organization and symmetry breaking in intestinal organoid development. Nature 569: 66–72

Sheu JJ-C, Guan B, Tsai F-J, Hsiao EY-T, Chen C-M, Seruca R, Wang T-L & Shih I-M (2012) Mutant BRAF Induces DNA Strand Breaks, Activates DNA Damage Response Pathway, and Up-Regulates Glucose Transporter-1 in Nontransformed Epithelial Cells. Am J Pathol 180: 1179–1188

Shimokawa M, Ohta Y, Nishikori S, Matano M, Takano A, Fujii M, Date S, Sugimoto S, Kanai T & Sato T (2017) Visualization and targeting of LGR5+ human colon cancer stem cells. Nature 545: 187–192

Smillie CS, Biton M, Ordovas-Montanes J, Sullivan KM, Burgin G, Graham DB, Herbst RH, Rogel N, Slyper M, Waldman J, et al (2019) Intra- and Inter-cellular Rewiring of the Human Colon during Ulcerative Colitis. Cell 178: 714–730.e22

Stuart T, Butler A, Hoffman P, Hafemeister C, Papalexi E, Mauck WM, Hao Y, Stoeckius M, Smibert P & Satija R (2019) Comprehensive Integration of Single-Cell Data. Cell 177: 1888–1902.e21

Sun C, Fang Y, Labrie M, Li X & Mills GB (2020) Systems approach to rational combination therapy: PARP inhibitors. Biochem Soc Trans 48: 1101–1108

Tong K, Pellón-Cárdenas O, Sirihorachai VR, Warder BN, Kothari OA, Perekatt AO, Fokas EE, Fullem RL, Zhou A, Thackray JK, et al (2017) Degree of Tissue Differentiation Dictates Susceptibility to BRAF-Driven Colorectal Cancer. Cell Rep 21: 3833–3845

Vermeulen L, De Sousa E Melo F, van der Heijden M, Cameron K, de Jong JH, Borovski T, Tuynman JB, Todaro M, Merz C, Rodermond H, et al (2010) Wnt activity defines colon cancer stem cells and is regulated by the microenvironment. Nat Cell Biol 12: 468–476

van de Wetering M, Sancho E, Verweij C, de Lau W, Oving I, Hurlstone A, van der Horn K, Batlle E, Coudreuse D, Haramis AP, et al (2002) The beta-catenin/TCF-4 complex imposes a crypt progenitor phenotype on colorectal cancer cells. Cell 111: 241–250

Wolf FA, Angerer P & Theis FJ (2018) SCANPY: large-scale single-cell gene expression data analysis. Genome Biol 19: 15

Woolston A, Khan K, Spain G, Barber LJ, Griffiths B, Gonzalez-Exposito R, Hornsteiner L, Punta M, Patil Y, Newey A, et al (2019) Genomic and Transcriptomic Determinants of Therapy Resistance and Immune Landscape Evolution during Anti-EGFR Treatment in Colorectal Cancer. Cancer Cell 36: 35–50.e9

Zanconato F, Cordenonsi M & Piccolo S (2016) YAP/TAZ at the Roots of Cancer. Cancer Cell 29: 783–803

Zhan T, Ambrosi G, Wandmacher AM, Rauscher B, Betge J, Rindtorff N, Häussler RS, Hinsenkamp I, Bamberg L, Hessling B, et al (2019) MEK inhibitors activate Wnt signalling and induce stem cell plasticity in colorectal cancer. Nat Commun 10: 2197

Zhang L, Li Z, Skrzypczynska KM, Fang Q, Zhang W, O’Brien SA, He Y, Wang L, Zhang Q, Kim A, et al (2020) Single-Cell Analyses Inform Mechanisms of Myeloid-Targeted Therapies in Colon Cancer. Cell 181: 442–459.e29

