## Supplementary Figures S1-S8 for "Mitogen-activated protein kinase activity drives cell trajectories in colorectal cancer"

Florian Uhlitz[1,2,3,*], Philip Bischoff[1,4,*], Stefan Peidli[1,2,*], Anja Sieber[1,2], Benedikt Obermayer[5], Eric Blanc[5], Alexandra Trinks[1,4], Mareen Lüthen[1,3], Yana Ruchiy[1], Thomas Sell[1,2], Soulafa Mamlouk[1,3], Roberto Arsie[6], Tzu-Ting Wei[6], Kathleen Klotz-Noack[1,7], Roland F Schwarz[3,6], Birgit Sawitzki[7], Carsten Kamphues[3,8], Dieter Beule[5], Markus Landthaler[4,6], Christine Sers[1,3,4], David Horst[1,3,4], Nils Blüthgen[1,2,3,4,#] and Markus Morkel[1,3,4,#]

1 Charité Universitätsmedizin Berlin, Corporate Member of Freie Universität Berlin, Humboldt-Universität zu Berlin, and Berlin Institute of Health, Institute of Pathology, Charitéplatz 1, 10117 Berlin, Germany

2 IRI Life Sciences, Humboldt University of Berlin, Philippstrasse 13, 10115 Berlin, Germany

3 German Cancer Consortium (DKTK) Partner Site Berlin, German Cancer Research Center (DKFZ), 69120 Heidelberg, Germany

4 Berlin Institute of Health (BIH), Anna-Louisa-Karsch-Straße 2, 10178 Berlin, Germany

5 Charité Universitätsmedizin Berlin, Corporate Member of Freie Universität Berlin, Humboldt-Universität zu Berlin, and Berlin Institute of Health, Core Unit Bioinformatics (CUBI), Charitéplatz 1, 10117 Berlin, Germany

6 Berlin Institute for Medical Systems Biology (BIMSB), Max Delbrück Center for Molecular Medicine, Hannoversche Strasse 28, 10117 Berlin, Germany

7 Charité Universitätsmedizin Berlin, Corporate Member of Freie Universität Berlin, Humboldt-Universität zu Berlin, and Berlin Institute of Health, Institute of Medical Immunology, Augustenburger Platz 1, 13353 Berlin

8 Charité Universitätsmedizin Berlin, Corporate Member of Freie Universität Berlin, Humboldt-Universität zu Berlin, and Berlin Institute of Health, Department of Surgery, Hindenburgdamm 30, 12203 Berlin

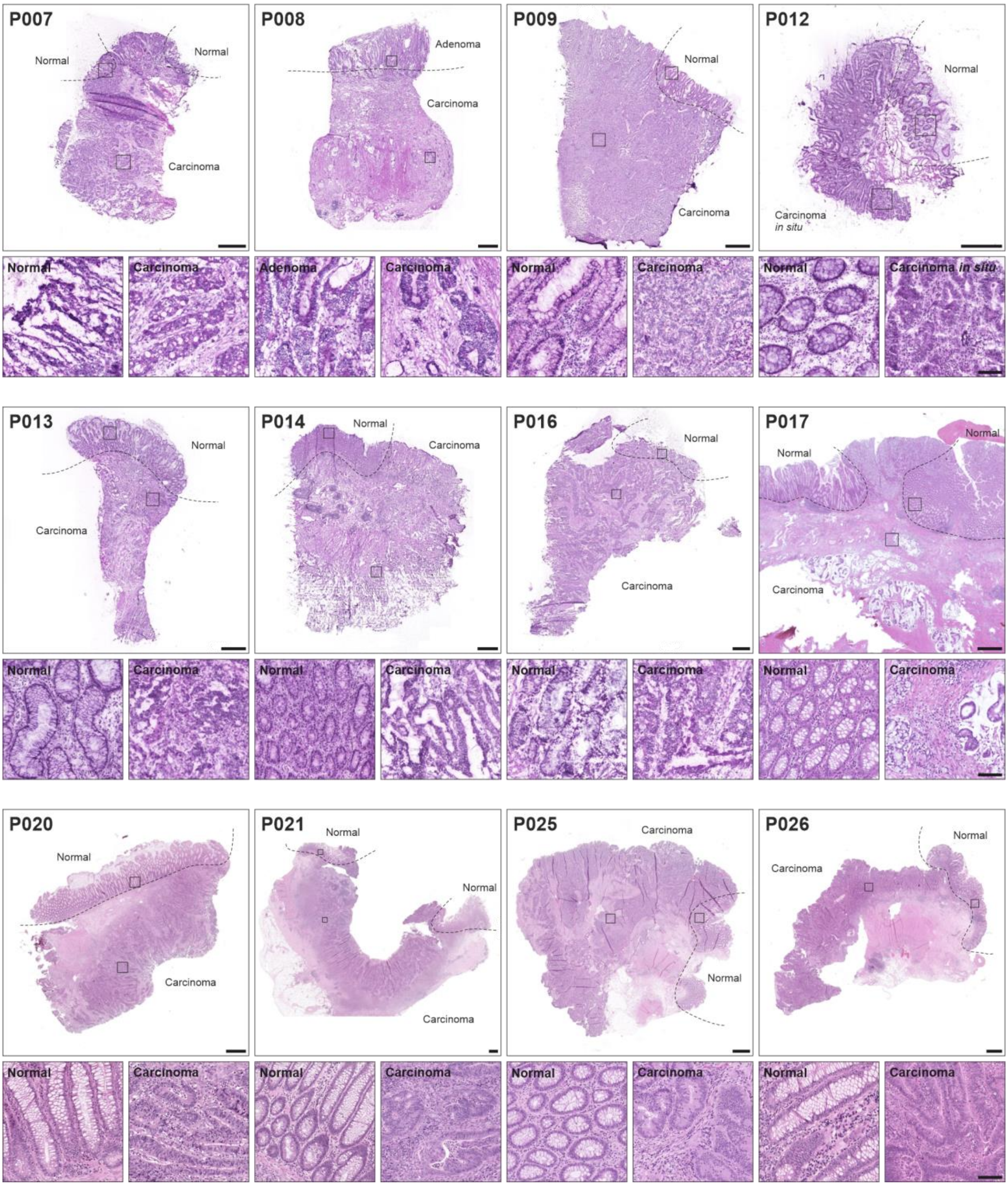


**Supplementary Figure 1: HE stainings of tumor samples used in the study.** All sections were done using adjacent fresh frozen material. Carcinoma, adenoma and normal tissue areas are indicated and magnified to the right of each panel.

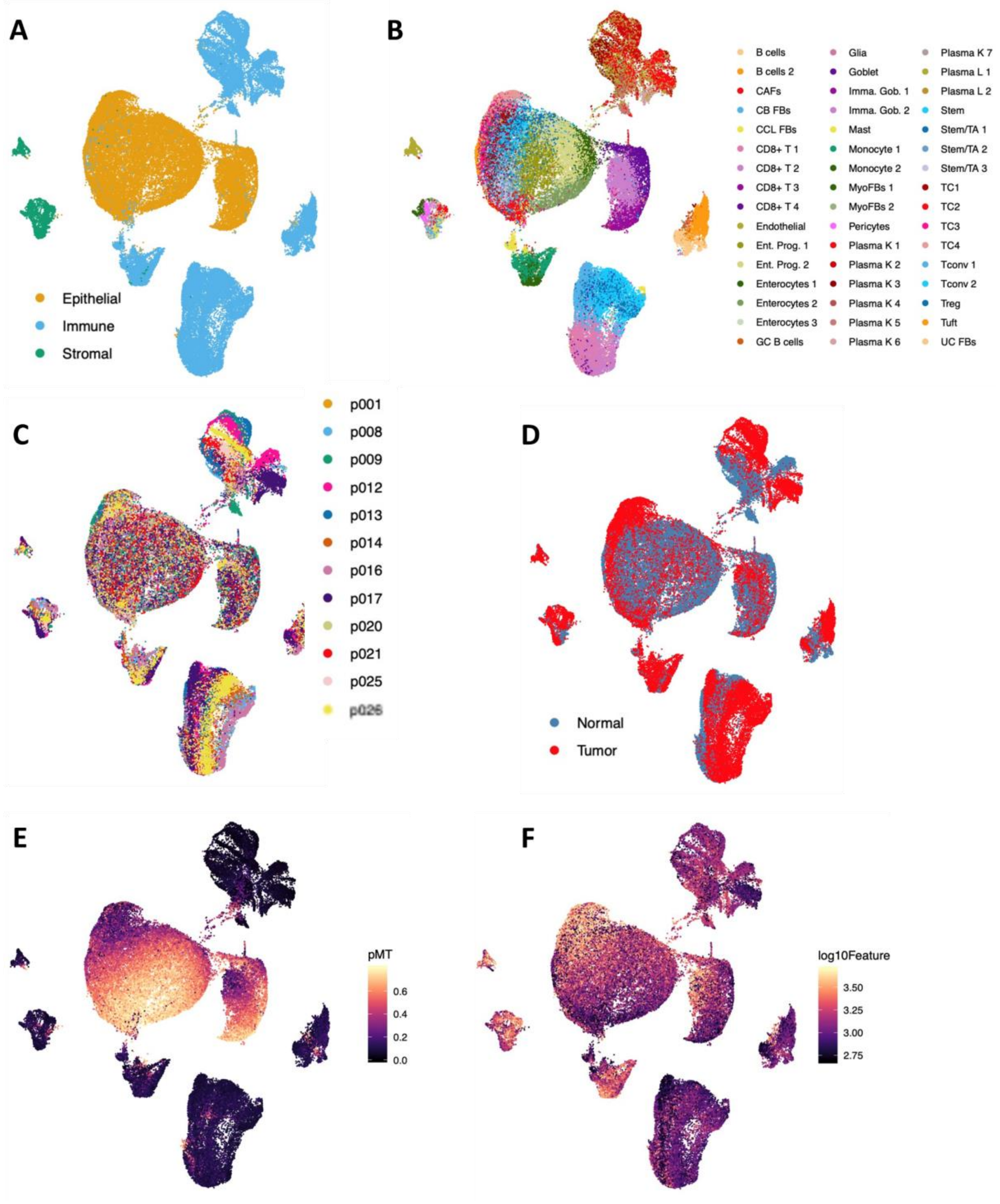


**Supplementary Figure 2: Initial clustering and single cell quality controls. A-F** UMAP of all single cell transcriptomes**. A** color coded by cluster assignment to epithelial, immune or stromal. **B** Final cell type assignment **C** color-coded by patinet. **D** color coded by tissue of origin. **E** color coded by percentage of mitochondrial reads. It is of note that the quality of epithelial cell transcriptomes is lower, in line with previous published studies. **F** color-coded by number of features (genes) per cell. See material and methods for further details.

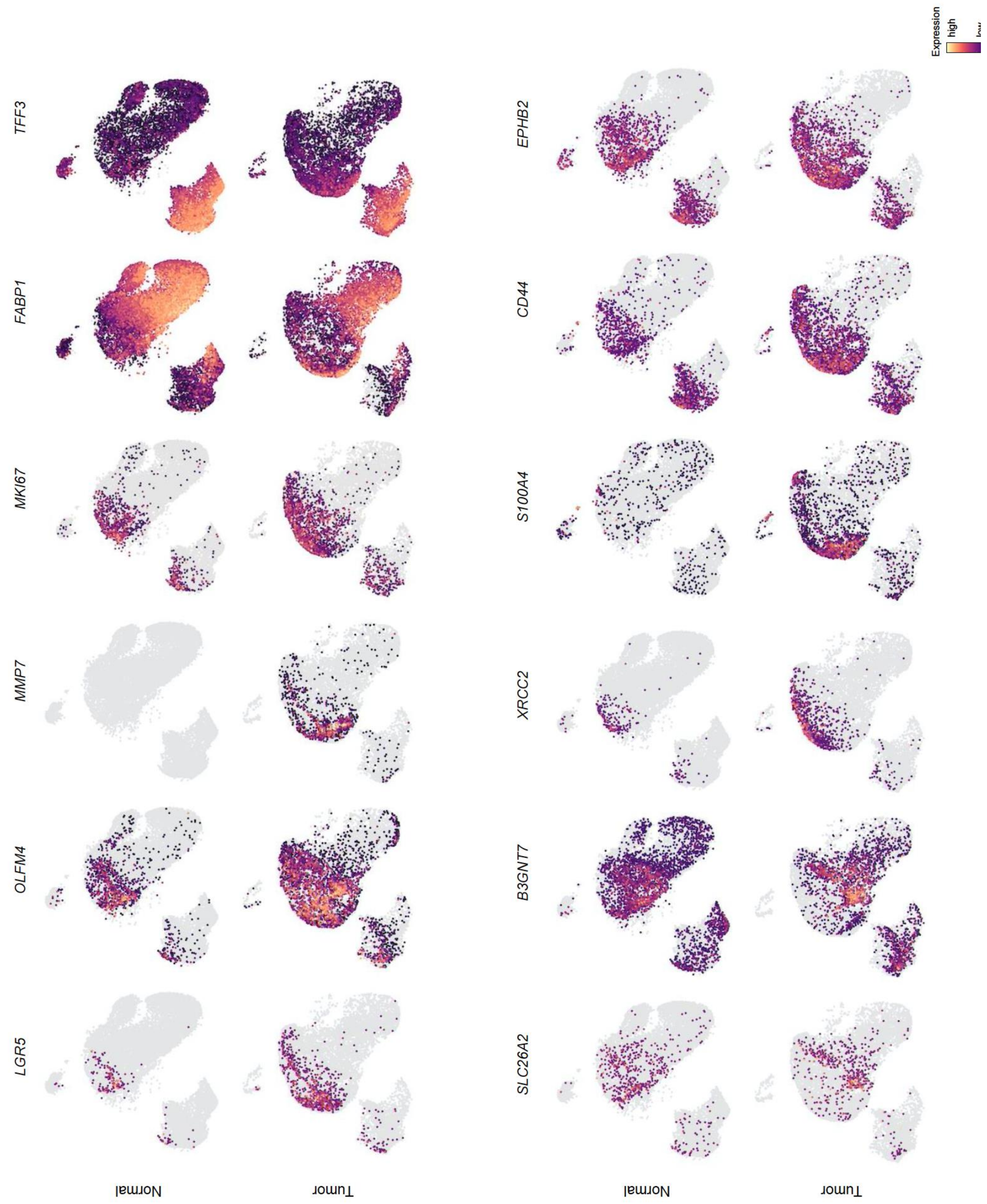


**Supplementary Figure 3: Expression of selected epithelial marker genes**. *LGR5* and *OLFM4* are colon stem cell markers; MMP7 is not expressed in the non-cancerous colon, but expressed in CRC; MKI67 is a proliferation marker; FABP1 and TFF3 are absorptive and secretory (goblet cell) differentiation markers, respectively; *SLC26A2* is enriched in enterocyte progenitors; *B3GNT7* is enriched in stem cells; *XRCC2* is a replication-stress related gene and a marker for TC1; *S100A4* is enriched in TC2-4; *CD44* is a stem cell marker enriched in TC1 and TC4; *EPHB2* is an intestinal crypt base marker enriched in TC1.

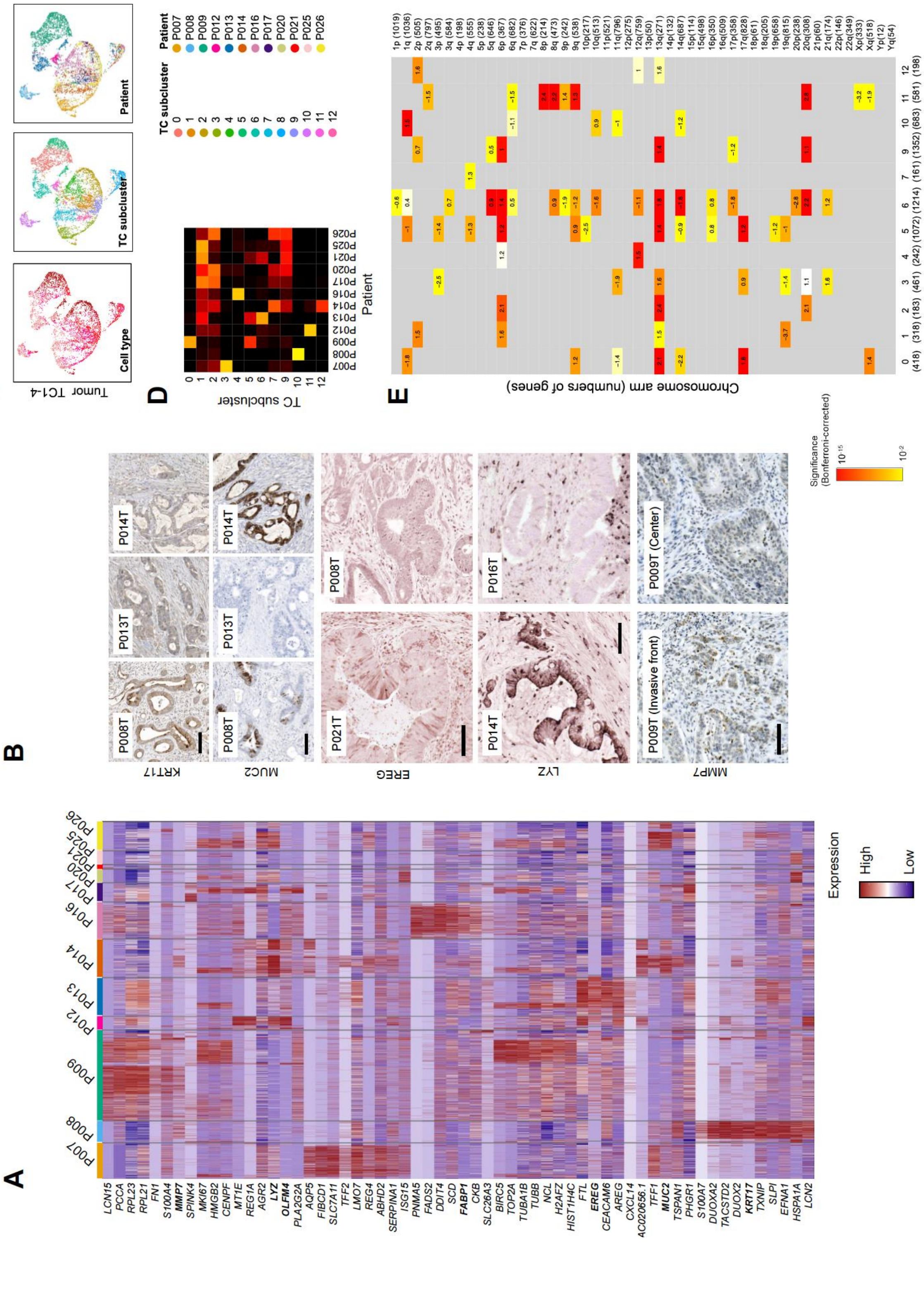


**Supplementary Figure 4: Analysis of patient-specific gene expression patterns. A** Heatmap of patient-specific gene expression. Genes validated by immunohistochemistry or immunofluorescence are given in bold. **B** Validation of patient-specific protein patterns, as indicated. **C** Subclustering of TC1-4 to reveal patient-specific components. **D** Heatmap showing relationship between TC1-4 subclusters and patients **E** overrepresentation of TC subcluster genes in genomic regions. Observed versus expected gene numbers were calculated using a Bonferroni-corrected hypergeometric distribution test, and human genome GRCh38 assembly gene numbers per chromosome arm as reference.

A

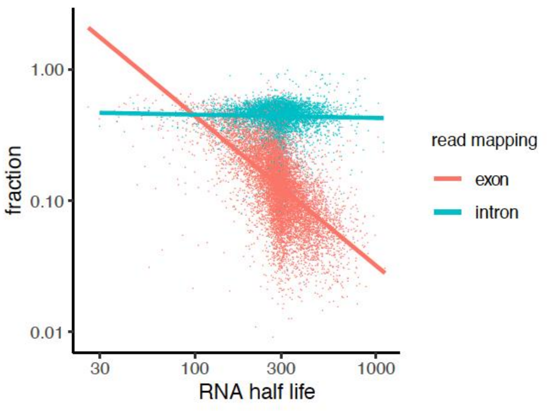


B

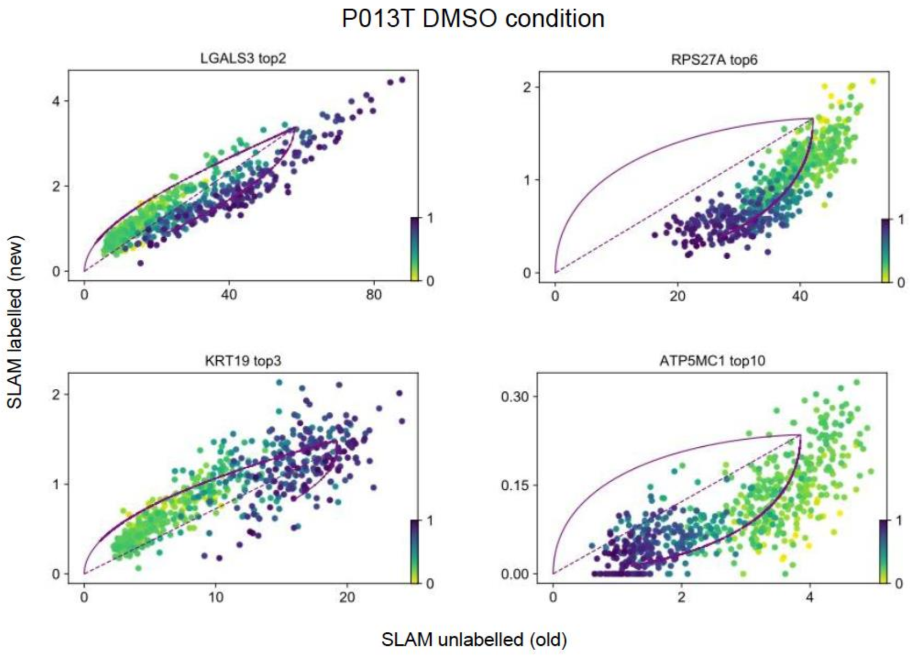


**Supplementary Figure 5: scSLAM-seq quality controls. A** Fraction of labeled reads for each transcript is function of the transcript's half-life (Caroline C. Friedel, Lars Dölken, Zsolt Ruzsics, Ulrich Koszinowski, Ralf Zimmer, Conserved principles of mammalian transcriptional regulation revealed by RNA half-life, Nucleic Acids Research, vol. 37, no. 17, pp. e115, 2009.) for reads covering introns (blue) or exons (red). This pattern is in agreement with successful labelling. **B** Phase plots for selected genes with ON or OFF dynamics. To the left: Differentiation markers *LGAL3* and *KRT19* show prototypical ON dynamics (increasing expression for late time points in the developmental trajectory). Ribosomal protein *RPS27A* and metabolic regulator *ATP5MC1* show prototypical OFF dynamics (high expression in early phases of the developmental trajectory, decreasing expression at late points of the cell developmental trajectory)

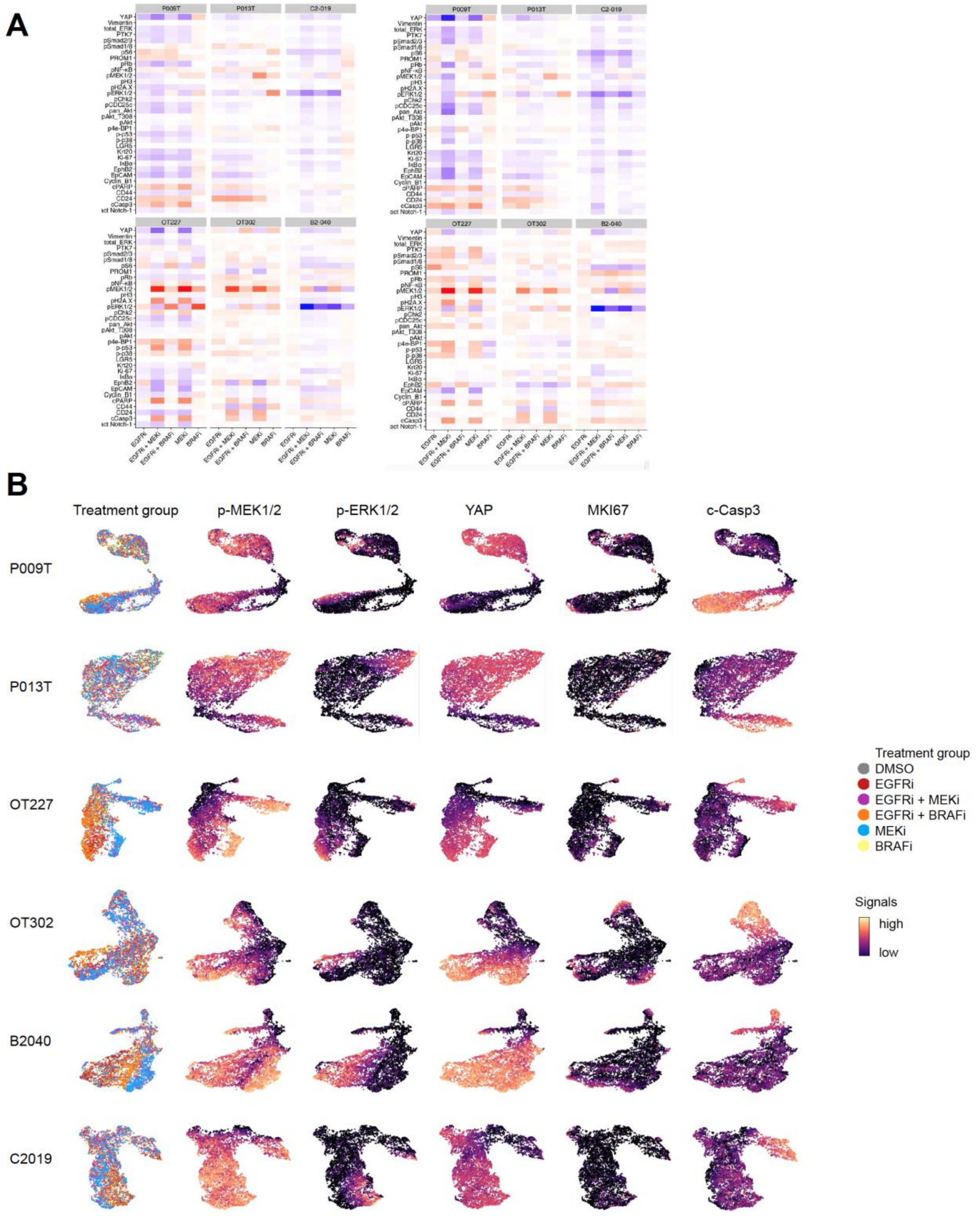


**Supplementary Figure 6: CyTOF quality controls and UMAPs. A** Heatmaps of independent replicate CyTOF runs. Heatmaps in main figure 5C were done from replicate 1. **B** CyTOF data of key signal transducers shown as UMAP.

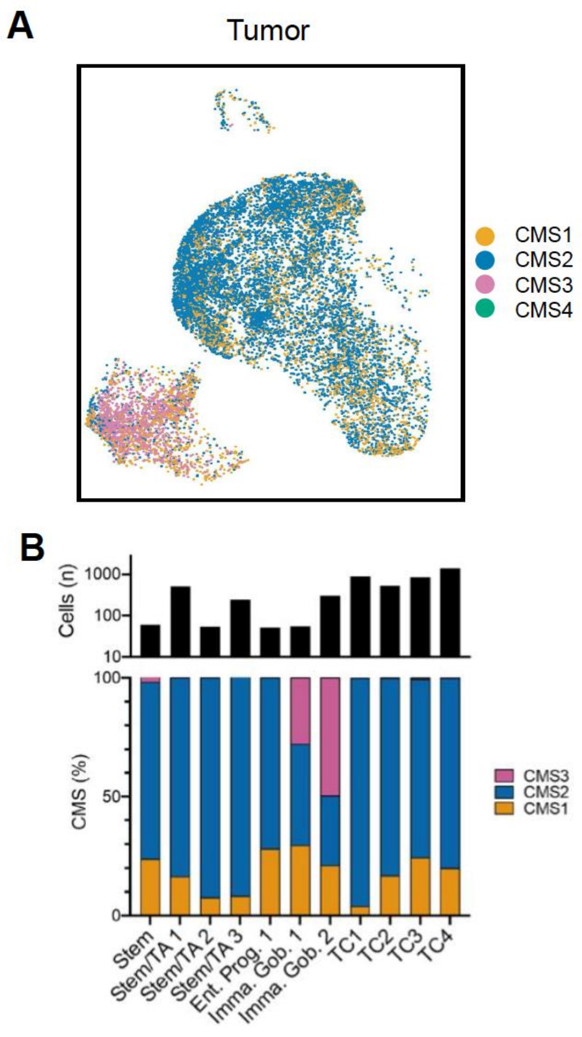


**Supplementary Figure 7: Assignment of CMS subtypes in tumor single cell sequencing data. A.** UMAPs of tumor tissue single cell data, as in main Fig. 1C, color-coded by CMS classifier. For exact localization of cell type clusters, see main Fig. 1C. **B** Cell type distribution and CMS classification for SCN-aberrant tumor cells. Top: Numbers of SCN-aberrant tumor cells per cluster. Only clusters with >50 SCNA cells are given. Below: CMS classification for SCN-aberrant cells, per cluster.

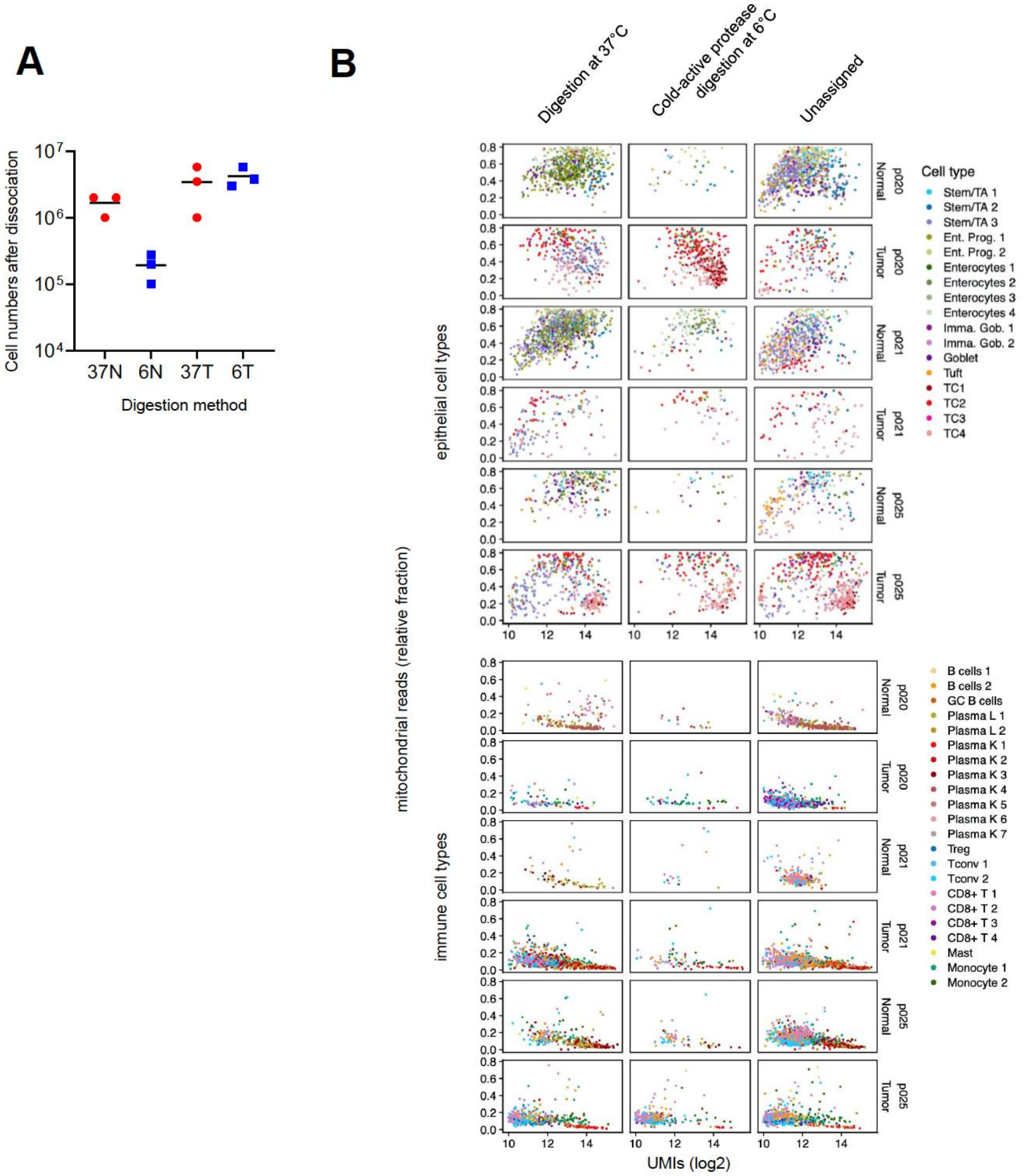


**Supplementary Figure 8: Test of alternative dissociation using cold active protease.** A cell type assignment of replicate samples digested at 37°C versus 6°C. **B** Qualities of epithelial and immune transcriptomes of replicate samples digested at 37°C versus 6°C for three patients P020, P021 and P025. Plots show fraction of mitochondrial reads versus unique molecule identifiers, color coded by assigned cell type. Replicate digestions were sequenced in the same library and were barcoded by BD sample tags. As the digestion interfered with sample tagging, many cells could not be assigned to the one or other digestion procedure, and thus remain unassigned (right column).
